# Comparative analysis reveals the long-term co-evolutionary history of parvoviruses and vertebrates

**DOI:** 10.1101/2021.10.25.465781

**Authors:** Matthew A. Campbell, Shannon Loncar, Robert Kotin, Robert J. Gifford

## Abstract

Parvoviruses (family *Parvoviridae*) are small DNA viruses that cause numerous diseases of medical, veterinary, and agricultural significance and have important applications in gene and anticancer therapy. DNA sequences derived from ancient parvoviruses are common in animal genomes and analysis of these *endogenous parvoviral elements* (EPVs) has demonstrated that the family, which includes twelve vertebrate-specific genera, arose in the distant evolutionary past. So far, however, such ‘paleovirological’ analysis has only provided glimpses into biology of parvoviruses and their long-term evolutionary interactions with hosts. Here, we comprehensively map EPV diversity in 752 published vertebrate genomes, revealing defining aspects of ecology and evolution within individual parvovirus genera. We identify 364 distinct EPV sequences and show these represent ∼200 unique germline incorporation events, involving at least five distinct parvovirus genera, that took place at points throughout the Cenozoic Era. We use the spatiotemporal and host range calibrations provided by these sequences to infer defining aspects of long-term evolution within individual parvovirus genera, including mammalian vicariance for genus *Protoparvovirus*, and inter-class transmission for genus *Dependoparvovirus*. Moreover, our findings support a model of virus evolution in which the long-term co-circulation of multiple parvovirus genera in vertebrates reflects the adaptation of each viral genus to fill a distinct ecological niche. Our discovery that parvovirus diversity can be understood in terms of genus-specific adaptations acquired over millions of years has important implications for their development as therapeutic tools - we show that these endeavours can now be approached from a rational foundation based on comparative evolutionary analysis. To support this, we published our data in the form of an open, extensible, and cross-platform database designed to facilitate the wider utilisation of evolution-related domain knowledge in parvovirus research.

## INTRODUCTION

Parvoviruses (family *Parvoviridae*) are a diverse group of small, non-enveloped DNA viruses that infect a broad and phylogenetically diverse range of animal species [1, 2]. The family includes numerous important pathogens of humans and domesticated species, including erythroparvovirus B19 (fifth disease) [3], carnivore protoparvovirus 1 (canine parvovirus) [4] and carnivore amdoparvovirus 1 (Aleutian mink disease; AMDV) [5]. Parvoviruses are also being developed as next-generation therapeutic tools: adeno-associated virus (AAV) has been successfully adapted as a gene therapy vector, and other parvoviruses are leading candidates for the further development as human gene therapy vectors [6, 7], while rodent protoparvoviruses (RoPVs) are promising anticancer agents that show natural oncotropism and oncolytic properties [8-10].

Parvoviruses have highly robust, icosahedral capsids (T=1) that contain a linear, single-stranded DNA genome typically ∼5 kilobases (kb) in length. Their genomes are very compact and generally exhibit the same basic genetic organization comprising two major gene cassettes, one (Rep/NS) that encodes the non-structural replication proteins, and another (Cap/VP) that encodes the structural coat proteins of the virion [11]. However, some genera contain additional open reading frames (ORFs) adjacent to these genes or overlapping them in alternative reading frames. Parvovirus genomes are flanked at the 3’ and 5’ ends by palindromic inverted terminal repeat (ITR) or ‘telomere’ sequences that are the only *cis* elements required for replication.

To understand the biology of parvoviruses it is helpful to have knowledge of their evolutionary history, because this can provide crucial insights into host-virus relationships and the biological basis of virus adaptations. Recent years have seen many important advances in understanding of parvovirus evolution and diversity, driven primarily by dramatic increases in the availability of DNA sequence data and investments in developing parvovirus-based therapeutics. A diverse range of novel parvovirus species have been described and the taxonomy of the family *Parvoviridae* has now been extensively re-organised to accommodate them [1]. In addition, progress in whole genome sequencing (WGS) has revealed that DNA sequences derived from parvoviruses are widespread within metazoan genomes [12-14]. These *endogenous parvoviral elements* (EPVs) arise when parvovirus infection of germline cells results in DNA becoming incorporated into chromosomal DNA so that integrated viral genes are then inherited as host alleles. EPV sequences can sometimes persist in the gene pool over many generations with the result that some are genetically ‘fixed’ (i.e., they reach a frequency of 100% in the species gene pool). These fixed EPV sequences have unique value to studies of parvovirus evolution because they provide a source of information about the long-term evolutionary history of interaction between parvoviruses and their hosts. Identification of orthologous EPV loci in multiple related host species demonstrates that integration occurred in their common ancestor, thereby providing robust minimum age estimates for parvovirus lineages, based on host species divergence times. Comparative studies have shown that many of the EPVs in vertebrate genomes derive from ancestral representatives of modern parvovirus genera, with some being derived from germline incorporation events that occurred >80 million years ago (Mya) [15-18].

The accumulation of genome sequence data from diverse, novel parvovirus species and EPVs presents unprecedented opportunities to investigate parvovirus biology using comparative approaches. Presently, however, there are two major obstacles to efficient use of parvovirus genome data in comparative investigations. Firstly, knowledge of the evolutionary timescale is still lacking - while EPVs have clearly established the ancient origins of certain parvovirus groups, a broader-scale analysis and a more detailed timeline are needed. Secondly, broad-scale, computational genomic analyses of viruses are often labour-intensive to implement, typically requiring the construction of complex data sets comprising sequences, multiple sequence alignments (MSAs) and phylogenies linked to other diverse kinds of data. High levels of sequence divergence greatly complicate the construction of sequence alignments and this problem is exacerbated when attempting to incorporate endogenous viral elements such as EPVs which tend (overall) to be neutrally evolving and hence are often highly degraded. A lack of tools and data standards currently prevents re-use and collaborative development of common resources for in-depth, comparative analysis of parvovirus genomes.

In this study, we directly address these obstacles – we catalogue EPV sequences in published WGS data and perform broad-scale comparative analysis of 752 published vertebrate genomes to recover 364 distinct EPV sequences representing at least 199 unique loci and involving at least five distinct parvovirus genera. We perform a broad-scale phylogenetic and genomic analysis of vertebrate parvoviruses, encompassing all known vertebrate parvovirus species as well as all vertebrate EPVs, and show that spatiotemporal and host range calibrations obtained from EPV sequences allow us to infer defining aspects of long-term evolution within individual parvovirus genera. Through broad-scale mapping of EPV diversity in vertebrates we establish a detailed picture of their long-term history of interaction with parvoviruses. In addition, we address issues of low reproducibility and re-use by publishing our data in the form of an open, extensible database “Parvovirus-GLUE”. This open, cross-platform resource utilizes a relational database framework to preserve the semantic links between the data items it contains (i.e., EPV and virus genome sequences, genome annotations, multiple sequence alignments, phylogenies, sequence-associated tabular data). Furthermore, by hosting Parvovirus-GLUE in an online version control system (GitHub), we establish a platform for the collaborative mapping of parvovirus/EPV diversity, and the wider utilisation of evolution-related domain knowledge in parvovirus research.

## RESULTS

### Creation of open resources for reproducible genomic analysis of parvoviruses

To facilitate greater reproducibility and reusability in comparative genomic analyses we previously developed GLUE (Genes Linked by Underlying Evolution), a bioinformatics software framework for the development and maintenance of ‘virus genome data resources’ [19]. Here, we used the GLUE framework to create Parvovirus-GLUE [20], an openly accessible online resource for comparative analysis of parvovirus genomes (**Fig. S1, Fig. S2**). Data items collated in Parvovirus-GLUE include: (i) a set of 135 reference genome sequences (**Table S1**) each representing a distinct parvovirus species and linked to isolate-associated data (isolate name, time and place of sampling, host species); (ii) a standardized set of 51 parvovirus genome features (**Table S2**); (iii) genome annotations specifying the coordinates of these genome features within reference genome sequences (**Table S3**); (iv) a set of MSAs constructed to represent distinct taxonomic levels within the family *Parvoviridae* (**Table 1, Fig. S3**). Standardised, reproducible comparative genomic analyses can be implemented by using GLUE’s command layer to coordinate interactions between the database and bioinformatics software tools (see **Methods**).

**Table 1.**
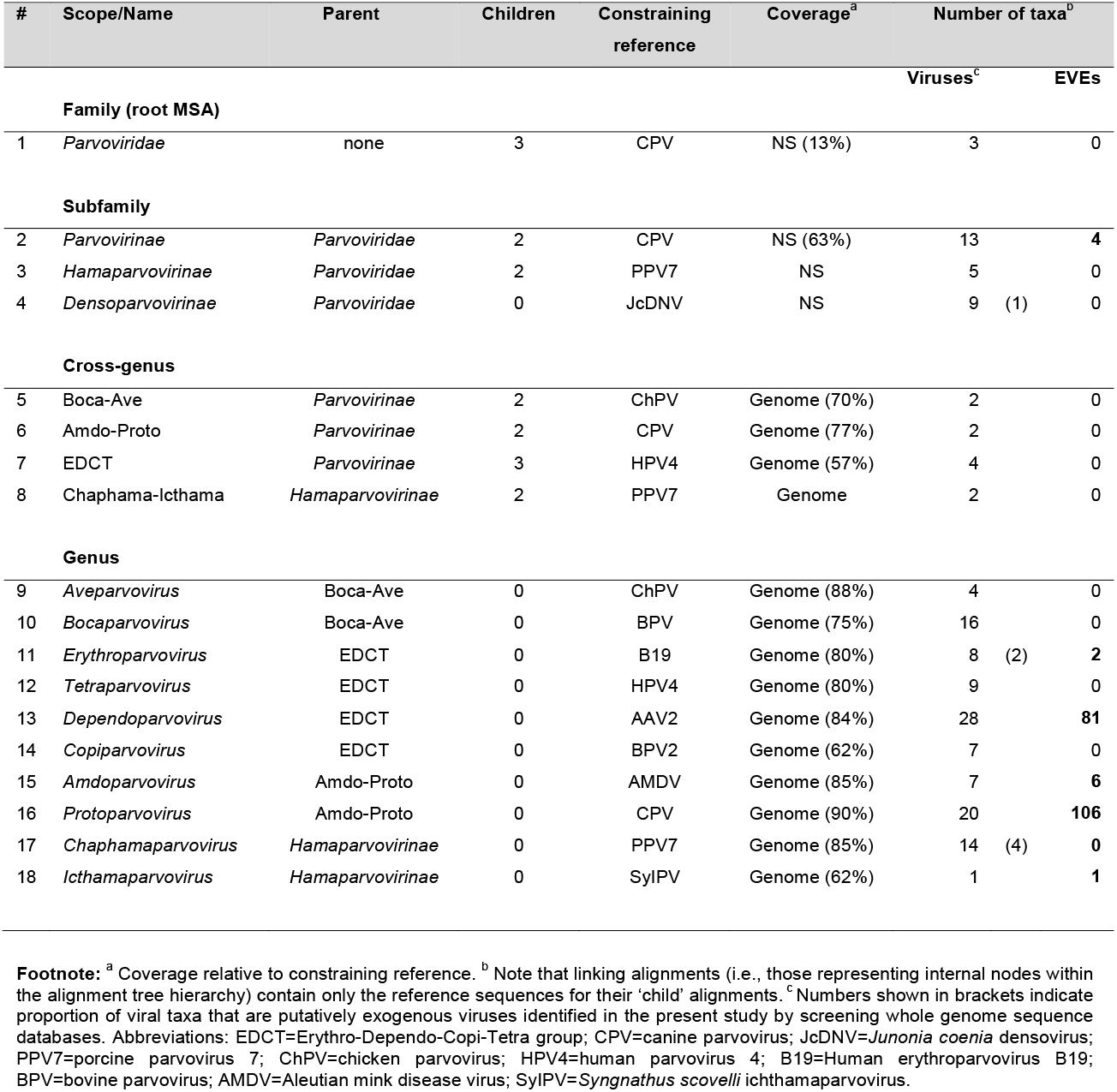
Summary of multiple sequence alignment (MSA) hierarchy constructed for the family *Parvoviridae*.

### Comprehensive mapping of endogenous parvoviral elements in published vertebrate genomes

To identify EPV loci in published vertebrate genomes, we performed systematic, similarity search-based *in silico* screening (see **Fig. S5**) of WGS data representing 752 vertebrate species. This led to the recovery a total of 595 EPV sequences (**Fig. 1**) which we resolved into a set of 199 distinct orthologous loci using comparative approaches (**Fig. 2**). We investigated the genomic regions flanking each putatively novel locus and identified flanking genes for most EPV loci (**Tables S4-S9**). Next, we compiled the robust, orthology-based minimum age calibrations we obtained from EPVs to generate an overview of parvovirus and vertebrate interaction over the past 100 My (**Fig. 3**). The oldest known EPV, a short VP-derived insert located in the *limbin* gene locus [15] - was not detected by the virus probes used in our initial screen (although it was detected when EPV-based probes were used). These findings imply that – as might be expected – our capacity to detect parvovirus EVEs is limited by their age. Notably, the limit of detection for parvoviruses seems to be ∼100 My which is similar to that found for bornaviruses (family *Bornaviridae*) which have RNA genomes, than found for reverse-transcribing DNA viruses (*Hepadnaviridae*) [21].

**Figure 1.**
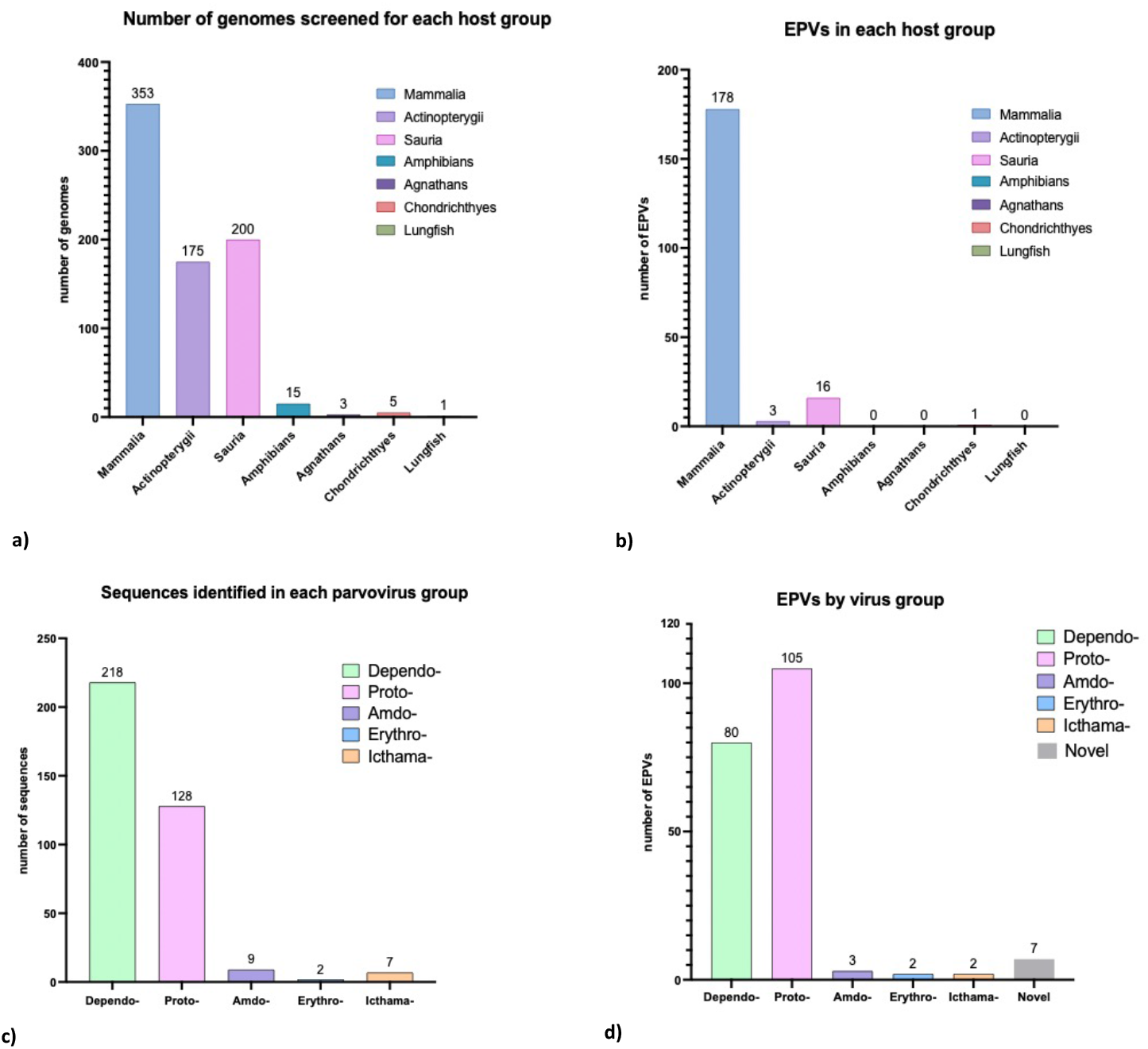
Summary of EPV diversity identified via *in silico* screening. **(a)** Number of genomes screened per host class. The number of screened genomes per host class is shown above the corresponding bar. Sauria is comprised of birds (144 genomes) and reptiles (56 genomes). 752 genomes were screened in total. **(b)** Number of sequences identified in each parvovirus group. The number of endogenous parvoviral element sequences identified in the screen is shown above each bar. 364 sequences were identified in total (excluding unclassified sequences). **(c)** Number of sequences identified in each parvovirus group. The number of endogenous parvoviral element sequences identified in the screen is shown above each bar. 364 sequences were identified in total (excluding unclassified sequences). Graphs were plotted with GraphPad Prism9.

**Figure 2.**
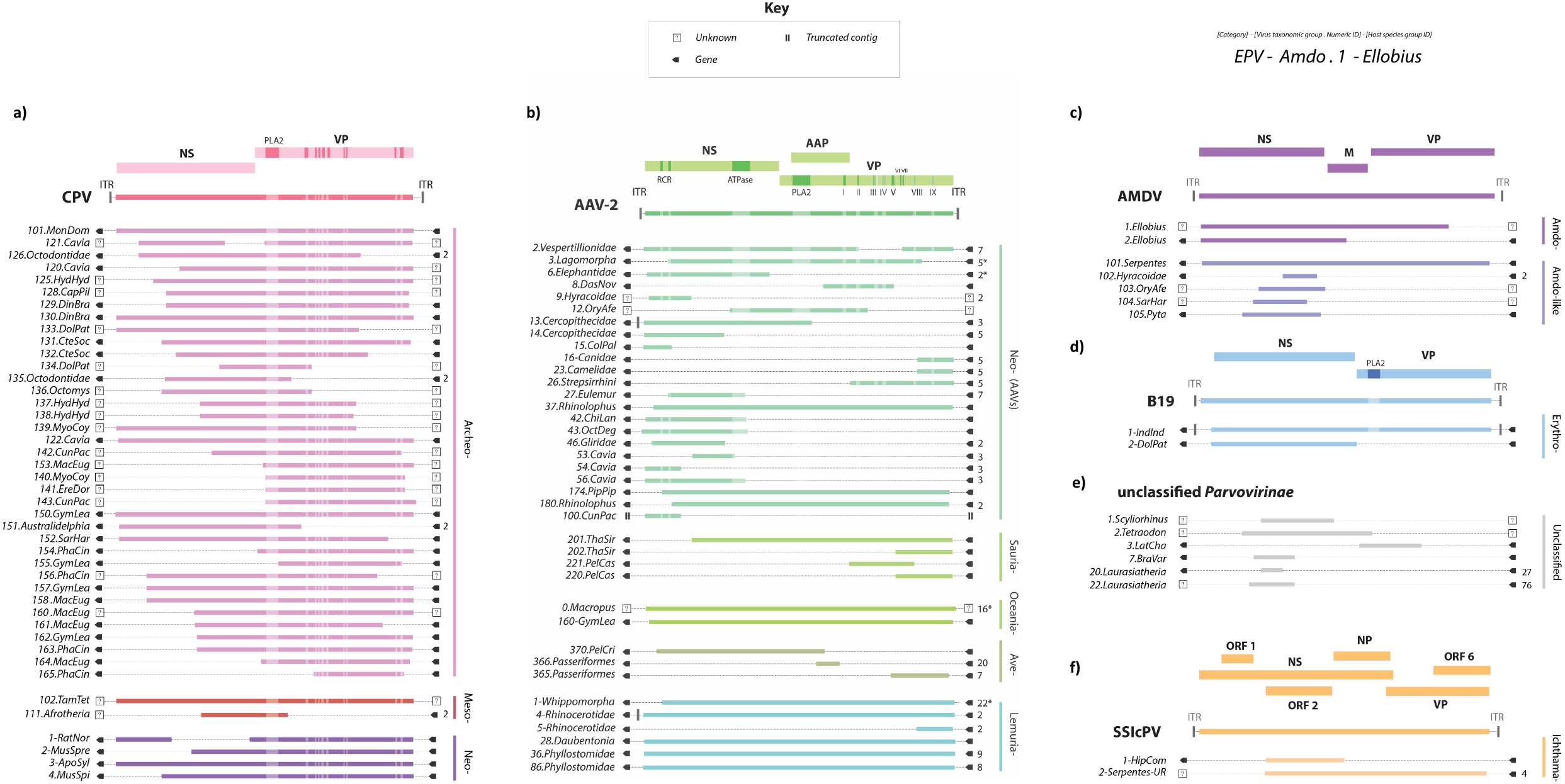
Genomic structures of unique EPV loci. **(a)** Protoparvovirus-derived EPV loci shown relative to the canine parvovirus (CPV) genome; **(b)** Dependoparvovirus-derived EPVs loci shown relative to the adeno-associated virus 2 (AAV-2) genome; **(c)** EPV loci derived from Amdoparvovirus-like viruses shown relative to the Aleutian mink disease (AMDV) genome; **(d)** Erythroparvovirus-derived loci shown relative to the parvovirus B19 genome; **(e)** EPVs derived from unclassified parvoviruses shown relative to a generic parvovirus genome. **(f)** Icthamaparvovirus-derived loci shown relative to *Syngnathus scovelli* parvovirus (SscPV). EPV locus identifiers are shown on the left. Solid bars to the right of each EPV set show taxonomic subgroupings below genus level. Numbers shown to the immediate right indicate a consensus and the number of orthologs used to create it. Boxes bounding EPV elements indicate either (i) the presence of an identified gene (see **Tables S5-S10**), (ii) an uncharacterised genomic flanking region, or (iii) a truncated contig sequence (see key). EPV locus identifiers use six letter abbreviations to indicate host species. **Abbreviations**: NS=non-structural protein; VP=capsid protein; ORF=open reading frame. ITR=inverted terminal repeat; PLA2=phospholipase A2 motif.

**Figure 3.**
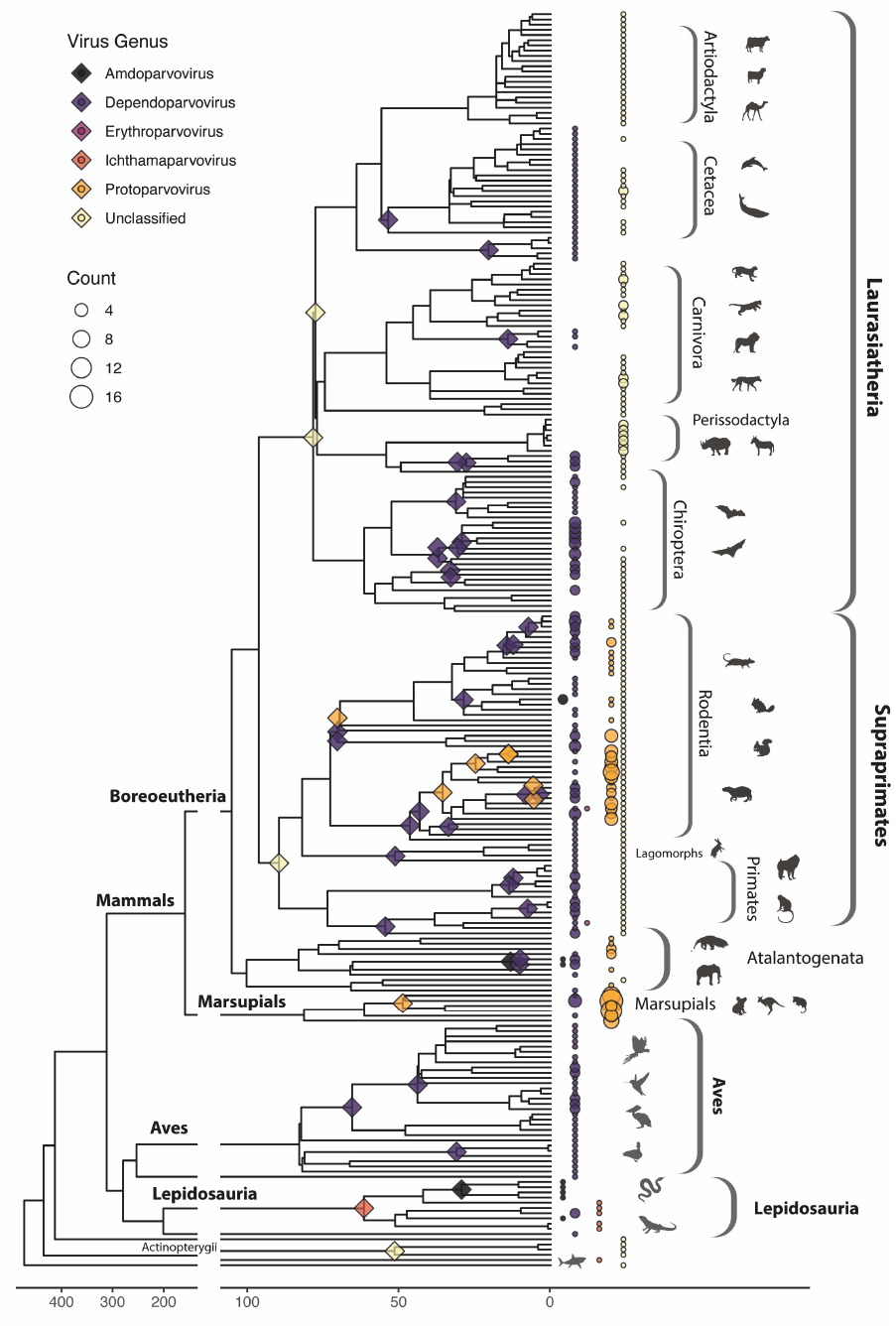
Incorporation of EPVs into the vertebrate germline. A time-calibrated evolutionary tree of vertebrate species examined in this study, illustrating the distribution of germline incorporation events over time. Colours indicate parvovirus genera as shown in the key. Diamonds on internal nodes indicate minimum age estimates for EPV loci endogenization (calculated for EPV loci found in >1 host species). Coloured circles adjacent to tree tips indicate the presence of EPVs in host taxa, with the diameter of the circle reflecting the number of EPVs identified (see count key). Brackets show taxonomic groups within vertebrates.

EPVs were identified in all major groups of terrestrial vertebrates except agnathans, crocodiles, and amphibians. Overall, however, they were found to occur much more frequently in mammalian genomes than in those of other vertebrate groups (**Table 2**). We used distance-based and phylogenetic approaches to taxonomically classify EPVs and found that they were predominantly derived from just two genera in subfamily *Parvovirinae -Protoparvovirus* and *Dependoparvovirus*. Other genera were represented, however, including the *Amdoparvovirus* and *Erythroparvovirus* genera of subfamily *Parvovirinae*, and the *Ichthamaparvovirus* genus of subfamily *Hamaparvovirinae* (**Fig. 1**). Other parvovirus genera known to infect vertebrates are conspicuously absent from the genomic fossil record: for example, no EPVs derived from the *Ave*-, *Boca*-, *Tetra-* and *Copiparvovirus* genera were identified.

**Table 2.** Incorporation of parvovirus DNA into the vertebrate germline.

| Parvovirus<br>genus | Host species group |  |  |  |  |  |  |  |  |  |
| --- | --- | --- | --- | --- | --- | --- | --- | --- | --- | --- |
|  | Chondrichthyes<br>*species=5* |  | Actinopterygii<br>*species =175* |  | Sauria<br>*species =200* |  | Mammalia<br>*species=353* |  | Vertebrata<br>*species =752\** |  |
|  | *loci* | *ratio* | *loci* | *ratio* | *loci* | *ratio* | *loci* | *ratio* | *loci* | *ratio* |
| *Ichthamaparvovirus* | 0 | - | 1 | 0.01 | 2 | 0.01 | 0 | - | 2 | 0.003 |
| *Erythroparvovirus* | 0 | - | 0 | - | 0 | - | 2 | 0.01 | 2 | 0.003 |
| *Amdoparvovirus* | 0 | - | 0 | - | 1 | 0.01 | 2 | 0.01 | 3 | 0.004 |
| *Dependoparvovirus* | 0 | - | 0 | - | 13 | 0.07 | 66 | 0.19 | 80 | 0.108 |
| *Protoparvovirus* | 0 | - | 0 | - | 0 | - | 105 | 0.3 | 105 | 0.142 |
| Novel clades | 1 | 0.2 | 2 | 0.01 | 0 | - | 3 | 0.01 | 7 | 0.009 |
| Totals | <u>1</u> | <u>0.2</u> | <u>3</u> | <u>0.02</u> | <u>16</u> | <u>0.08</u> | <u>178</u> | <u>0.5</u> | <u>199</u> | <u>0.263</u> |

**Footnote:** \*Agnathans (n=3), Amphibians (n=15), and Lungfish (*Latimeria chalumnae*) were also screened but results are not shown since no species in these groups were found to have EPVs. Ratio = unique loci / genomes screened.

While the majority of EPV loci identified in our study are either unambiguous members of contemporary parvovirus genera or members of closely related sister groups, seven loci could not be phylogenetically classified. Some were relatively short and old – e.g., unclassified *Parvovirinae* elements 20 and 22 in Laurasiatheria, and 23 in Supraprimates, all of which are genomic fragments that are >80 million years old. Longer, yet still unclassifiable EPVs were identified in lower vertebrate groups (e.g., lobe-finned fish). Since parvovirus diversity has yet to be thoroughly investigated in lower fish and other lower vertebrates, these sequences might represent members of unknown viral genera that currently infect these host groups.

### Novel exogenous virus species identified in silico via similarity-search based screening

Our screen of WGS data identified numerous sequences that appear to derive from contaminating viruses, including both parvoviruses and herpesviruses (**Table S11**). A putative betaherpesvirus was identified via the presence of a U94 homolog - U94 is a parvovirus-derived gene that is encoded by the *Roseolovirus* genus of betaherpesviruses (subfamily *Betaherpesvirinae*) (**Fig. S9a-b**). In addition to the chaphamaparvoviruses previously reported in WGS data [22] we identified a divergent densoparvovirus in WGS data of the annual killifish (*Austrofundulus limnaeus*) (**Fig. S9c**), and erythroparvovirus sequences in WGS data of the Masai giraffe (*Giraffa tippelskirchii*) and the striped hyena (*Hyaena hyaena*). Additionally, *ad hoc*, BLAST-based screening of mRNA databases identified protoparvovirus-like sequences in the gray bichir (*Polypterus senegalus*) (**Fig. S9d**).

### Long-term co-evolutionary interaction between vertebrates and subfamily Parvovirinae

We performed a comprehensive phylogenetic analysis of the subfamily *Parvovirinae*, which exclusively infects vertebrates (**Fig. 4, Fig. S7, Fig. S8**). Phylogenies reconstructed from aligned *rep* genes reveal three robustly supported sub-lineages each encompassing multiple genera, as follows: (1) “ETDC”: *Erythro*-, *Tetra*-, *Dependo*- and *Copiparvovirus* (**Fig.3a**); (2) “Ave-Boca”: *Ave*- and *Bocaparvovirus* (**Fig.3a**); (3) “Amdo-Proto”: *Amdo*- and *Protoparvovirus* (**Fig.3b**). The EDTC and “Amdo-Proto” clades are demonstrably ancient as they both include EPVs that were incorporated into the germline >80 Mya, and while the “Ave-Boca” lineage does not have fossil representatives, it is interesting to note that it comprises entirely distinct saurian (i.e., avian) lineages and would be >200My old if it arose prior to the Mammalia-Sauria split (**Table 3, Fig.3**). Similarly, we identified EPVs derived from the “Amdo-Proto” and “ETDC” lineages in basal vertebrates including lobe-finned fish (class Sarcopterygii) and sharks (class Chondrichthyes), suggesting that the origins of these groups may predate the deep divergences among terrestrial vertebrates (**Table 3, Fig.3**).

**Figure 4.**
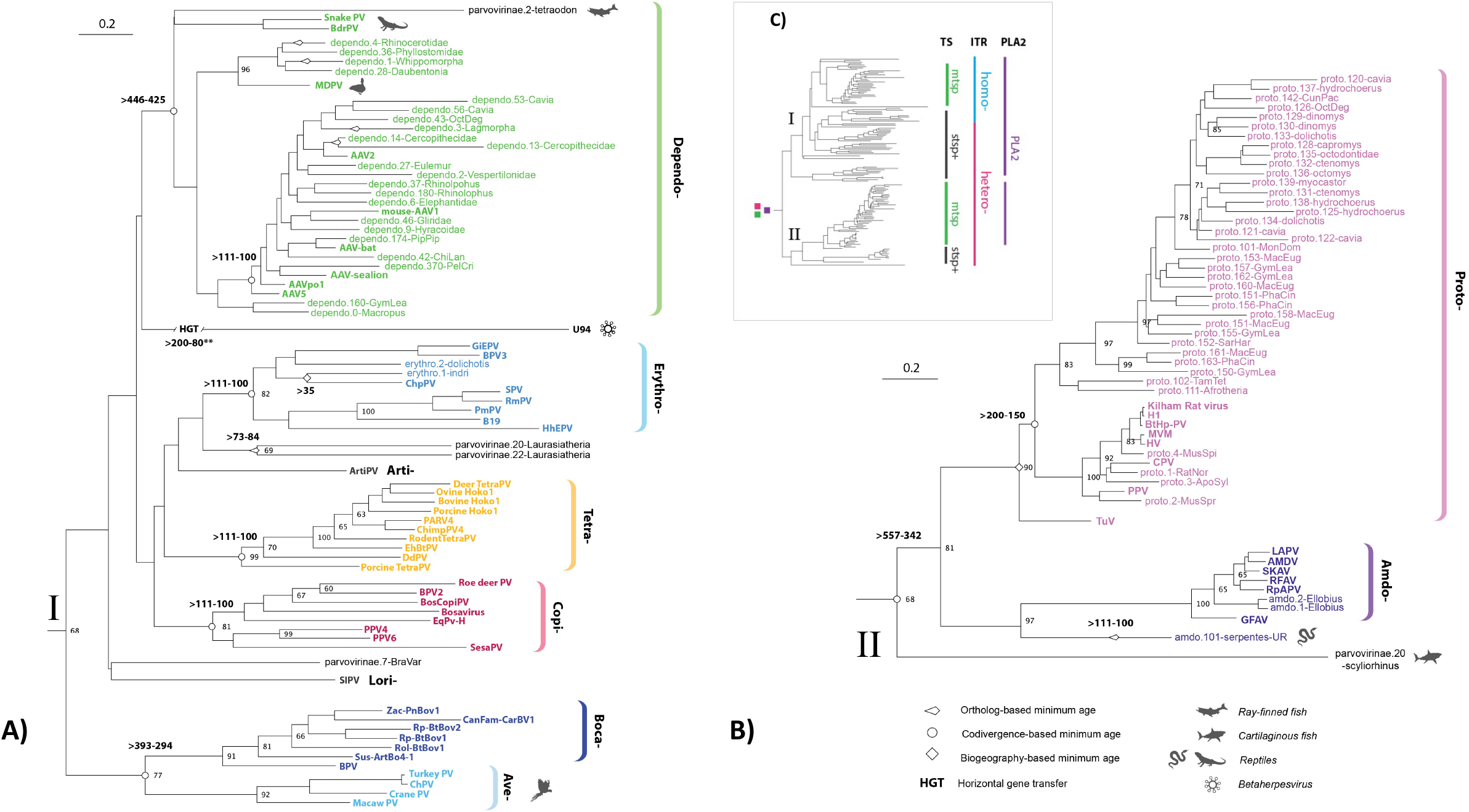
Evolution of subfamily *Parvoviridae*. A maximum likelihood phylogeny showing the reconstructed evolutionary relationships between contemporary parvoviruses and the ancient parvovirus species represented by endogenous parvoviral elements (EPVs). Panels (A) and (B) show a more detailed view of subclades (labelled I and II) within the phylogeny shown in panel (C). The complete phylogeny, which is midpoint rooted for display purposes, was reconstructed using a multiple sequence alignment spanning 270 amino acid residues in the Rep protein and the LG likelihood substitution model. Coloured brackets indicate the established parvovirus genera recognised by the International Committee for the Taxonomy of Viruses. Bootstrap support values (1000 replicates) are shown for deeper internal nodes only. Scale bars show evolutionary distance in substitutions per site. Taxa labels are coloured based on taxonomic grouping as indicated by brackets, unclassified taxa are shown in black. Viral taxa are shown in bold, while EPV taxa are show in regular text. EPV were assigned unique identifiers (IDs) constructed from three components. The first component is the classifier ‘EPV’. The second component comprises the name of the lowest level taxonomic group (i.e., species, genus, subfamily, or other clade) into which the element can be confidently placed by phylogenetic analysis and a numeric ID that uniquely identifies the insertion, separated by a period. The third component specifies the group of species in which the sequence is found. **Abbreviations**: PV=Parvovirus; HHV=Human herpesvirus; AAV=Adeno-associated virus; AMDV=Aleutian mink disease; CPV=canine parvovirus; BPV=bovine parvovirus; BrdPV=Bearded dragon parvovirus; MdPV=Muscovy duck parvovirus; SlPV=slow loris parvovirus. TS=transcription strategy; MTSP=Multiple transcriptional start positions; STSP+=single transcription start position, plus additional strategies; HOMO=homotelomeric; HETERO=heterotelomeric.

**Table 3.**
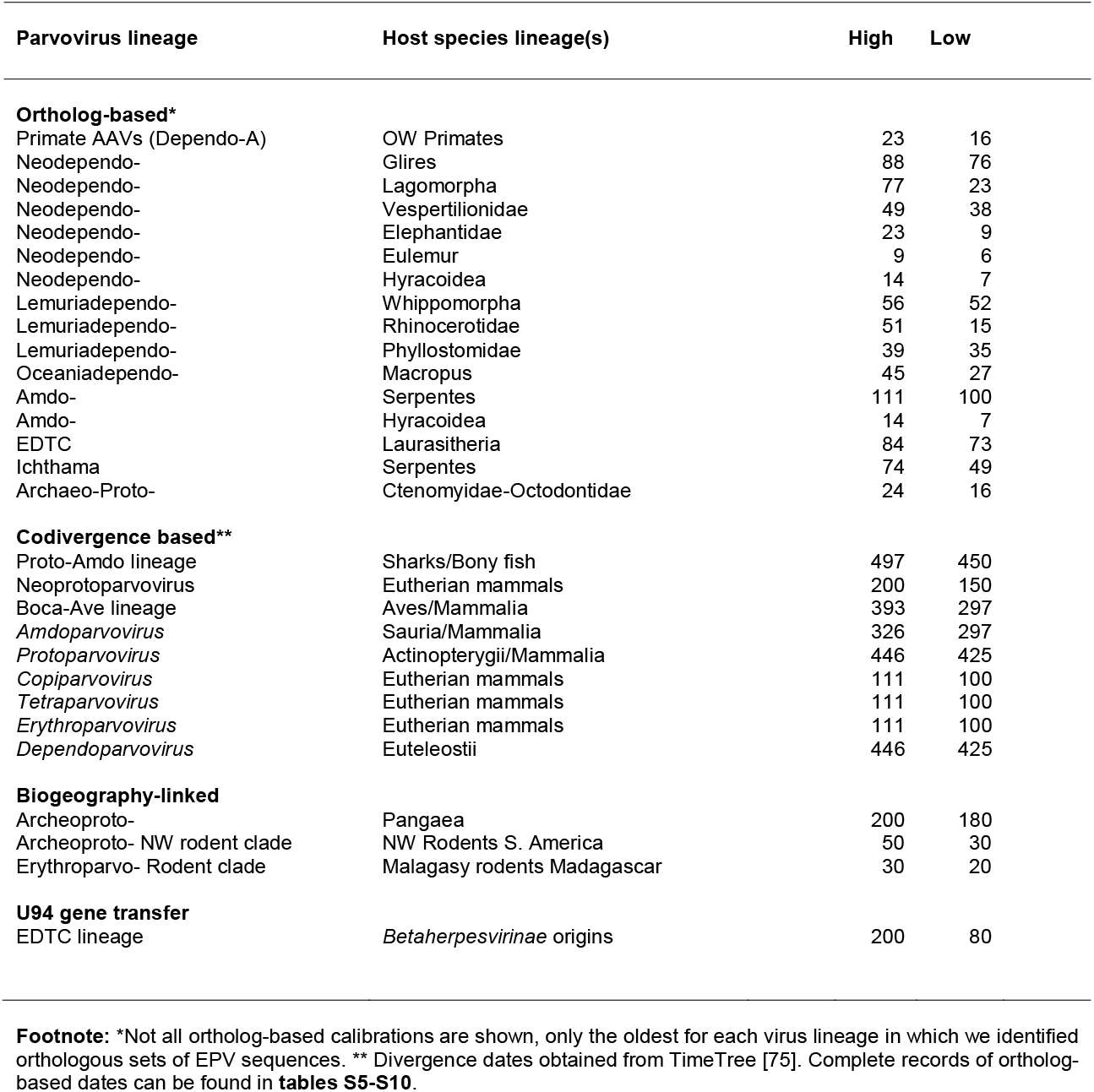
Dates and age estimates used to calibrate parvovirus evolution.

We examined the distribution of conserved genome features among *Parvovirinae* genera in relation to the *Parvovirinae* phylogeny (**Fig. 4c**). For example, the ‘telomeres’ that flank parvovirus genomes are heterotelomeric (asymmetrical) in some genera (*Amdo-, Proto-, Boca-*, and *Aveparvovirus*) whereas they are homotelomeric (symmetrical) in others [23]. Interestingly, the distribution of this trait across sub-lineages within the subfamily *Parvovirinae* suggests that the asymmetrical form (which is found across the “Amdo-Proto” and “Ave-Boca” sublineages) is more likely to be ancestral (**Fig. 4c**). Similarly, in all *Parvovirinae* groups, except genus *Amdoparvovirus*, the N-terminal region of VP1 (the largest of the capsid) contains a phospholipase A2 (PLA2) enzymatic domain that becomes exposed at the particle surface during cell entry and is required for escape from the endosomal compartments. Phylogenetic reconstructions indicate that this domain was present ancestrally and has been convergently lost in the *Aveparvovirus* and *Amdoparvovirus* genera (**Fig. 4c**) [2, 24].

*Parvovirinae* genera also show variation in their gene expression strategies through differential promoter usage and alternative splicing. Members of the *Proto-* and *Dependoparvovirus* genera use two to three separate transcriptional promoters whereas the *Amdo-, Erythro-*, and *Boca-* genera express all genes from a single promoter and use genus-specific read-through mechanisms to produce alternative transcripts [2, 11]. Interestingly, both the *Proto-* and *Dependoparvovirus* genera utilise the first of these expression strategies despite being relatively distantly related, suggesting that the use of separate promoters may be the ancestral strategy within the subfamily *Parvovirinae*, with other genera independently acquiring mechanisms to express multiple genes from a single promoter (**Fig 3c**).

### Horizontal gene transfer from the Parvovirinae to the Betaherpesvirinae

As discussed above, betaherpesviruses in the *Roseolovirus* genus, such as human herpesvirus 6 (HHV6), have acquired a homolog of the parvovirus *rep* gene, called U94. This gene presumably arose when co-infection of the same host cell led to parvovirus DNA being incorporated the ‘germline’ of a roseolovirus ancestor [25]. Our screen of vertebrate WGS data identified two homologs of the HHV6 U94 gene. One of these was embedded within an endogenous HHV6-derived sequence in the genome of the Philippine tarsier (*Carlito syrichta*), as reported previously [26], while the other was identified in WGS data of the stoat (*Mustela ermina*). The contig containing this sequence appears to correspond to a near complete betaherpesvirus genome (**Table S9**). Phylogenetic reconstructions indicate that U94 derives from a *Parvovirinae* sublineage (**Fig. 4a**). Furthermore, the presence of U94 homologs in primate [25, 26], rodent [27], and bat betaherpesviruses [28] as well as in the (presumably) carnivore betaherpesvirus described here, allow tentative estimation of when this gene arose. Given the demonstrable age of EPVs that fall robustly within the EDTC lineage, horizontal gene transfer likely occurred before the divergence of eutherian mammal orders (∼100 Mya) but after the estimated divergence of the three subfamilies (*Alpha*-, *Beta*-, *Gammaherpesvirinae*) of *Herpesviridae* (∼200) Mya [29] (**Table 3**). While U94 homologs have been retained in roseoloviruses demonstrates, the N-terminal domain of the protein is less well conserved across all four viruses examined here and has been lost entirely in one homolog (the r127 gene of rat cytomegalovirus), suggesting relatively strong diversifying selection (**Fig. S9**).

### Mammalian vicariance shaped the evolution of protoparvoviruses

Protoparvoviruses infect a broad range of mammalian hosts and cause a variety of conditions from subclinical to lethal disease [11]. The recovery of a rich fossil record for protoparvoviruses allowed us to examine how their evolution has been shaped by macroevolutionary processes impacting on mammals over the past 150-200 My, such as. continental drift [30]. Around 200 Mya, the supercontinent of Pangaea, then the sole landmass on the planet, began separating into two subcomponents (**Fig. 5b**). One (Laurasia) comprised Europe, North America and most of Asia, while the second (Gondwanaland) comprised Africa, South America, Australia, India, and Madagascar). Mammalian subpopulations were fragmented by these events, and then fragmented further as Gondwanaland separated into its component continents. The associated genetic isolation due to geographic separation (vicariance) drove the early diversification of major subgroups, including indigenous mammalian lineages in South America (xenarthans and marsupials), Australia (marsupials), and Africa (afrotherians). At points throughout the Cenozoic Era, placental mammal groups that evolved in Laurasia (boreoeutherians) expanded into other continental regions. For example, the ancestors of contemporary New World rodents (which include capybaras, chinchillas, and guinea pigs among many other, highly diversified species), are thought to have reached the South American continent ∼35 Mya [31].

**Figure 5.**
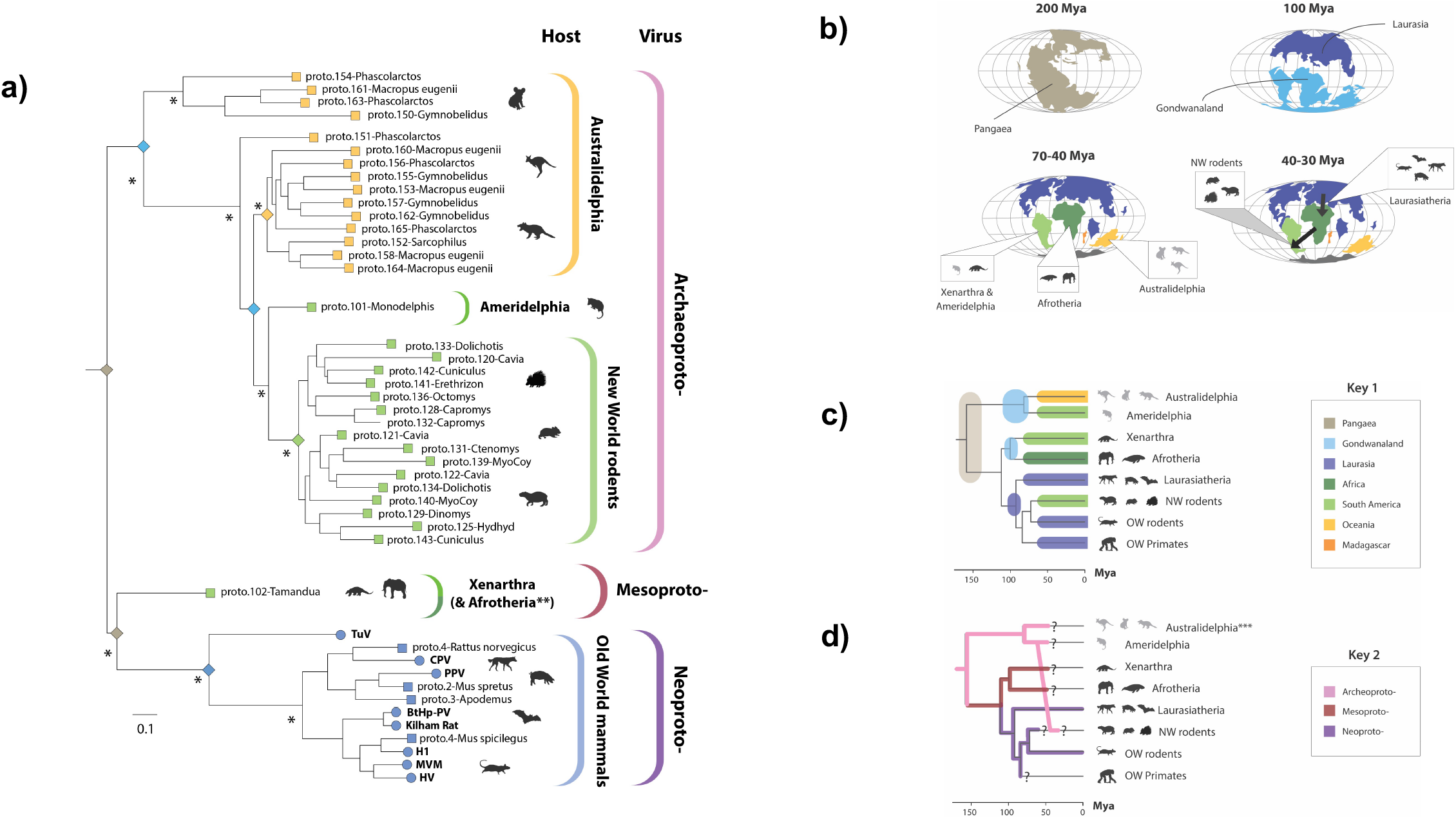
Protoparvovirus evolution has been shaped by mammalian vicariance. **(a)** Maximum likelihood-based phylogenetic reconstructions of evolutionary relationships between contemporary parvovirus species and the ancestral parvovirus species represented by endogenous parvoviral elements (EPVs). The phylogeny was constructed from a multiple sequence alignment spanning 712 amino acid residues in the Rep protein (substitution model=LG likelihood). The tree is midpoint rooted for display purposes. Asterisks indicate nodes with bootstrap support >70% (1000 replicates). The scale bar shows evolutionary distance in substitutions per site. Coloured brackets to the right indicate (i) subgroups within the *Protoparvovirus* genus (outer set of brackets) and (ii) the host range of each subgroup (inner set of brackets). Terminal nodes are represented by squares (EPVs) and circles (viruses) and are coloured based on the biogeographic distribution of the host species in which they were identified. Coloured diamonds on internal nodes show the inferred ancestral distribution of parvovirus ancestors, using colours that reflect the patterns of continental drift and associated mammalian vicariance shown in the maps in panel (b). **Evidence for the presence of the “Mesoprotoparvovirus” group in Afrotherians is presented in **Fig. 4. (b)** Mollweide projection maps showing how patterns of continental drift from 200-35 led to periods of biogeographic isolation for terrestrial mammals in Laurasia (Europe and Asia), South America, Australia Africa and Madagascar. The resulting vicariance is thought to have contributed to the diversification of mammals, reflected in the mammalian phylogeny as shown in in panel (c). Most placental mammals (including rodents, primates, ungulates and bats) evolved in Laurasia. However, these groups later expanded into other continents, and fossil evidence indicates that the ancestors of today’s “New World rodents” had arrived on the South American continent by∼35 million years ago (Mya), if not earlier. **(c)** A time-calibrated phylogeny of mammals with annotations indicating the biogeographic associations of the major taxonomic groups of contemporary mammals and ancestral mammalian groups, following panel (b) and key 1. **(d)** A time-calibrated phylogeny of mammals annotated to indicate the distribution of protoparvovirus subgroups among mammalian groups, following key 2. Question marks indicate where it is unknown whether viral counterparts of the lineages represented by EPVs still circulate among contemporary members of the host species groups in which they are found. **Abbreviations**: Mya = millions of years ago; NW=New World; NW=New World; OW=Old World; CPV=carnivore parvovirus type 1; PPV=porcine parvovirus; HV=Hamster parvovirus; TuV=Tusavirus.

We identified 121 protoparvovirus-related EPV sequences in mammals, which we estimate to represent at least 105 distinct germline incorporation events (**Table S1**). Several near full-length genomes were identified, and most elements spanned >50% of the genome (**Fig 1a**). Furthermore, protoparvovirus-like sequences identified in a fish transcriptome group basal to all mammalian protoviruses and protoparvovirus-like EPVs, and these sequences are all more closely related to one another than they are to amdoparvoviruses or divergent, amdoparvovirus-like EPVs (e.g., *Amdo*.*101-Serpentes*) [32] (**Fig 3b, Fig S9d**).

Phylogenies reveal previously unappreciated diversity within the *Protoparvovirus* genus: three major subclades are present, which we labelled “Archaeoproto-”, “Mesoproto-” and “Neoproto-” (**Fig 4a**). The *Archaeoprotoparvovirus* (ApPV) clade is comprised exclusively of EPVs and is highly represented in the genomes of Australian marsupials (Australidelphia), American marsupials (Ameridelphia) and New World rodents. The *Mesoprotoparvovirus* (MpPV) clade is also comprised exclusively of EPVs, and was sparsely represented in the EPV fossil record, only being detected in the genomes of basal placental mammal groups (Xenarthra and Afrotheria). Finally, the *Neoprotoparvovirus* (NpPV) clade contains all known contemporary protoparvoviruses and a small number of EPV elements derived from these viruses (**Fig. 5a**).

Protoparvovirus phylogenies strikingly reflect the impact of mammalian vicariance – and later migration – on their emergence and spread during the Cenozoic Era. As shown in **Fig. 5**, the protoparvovirus phylogeny can readily be mapped onto the phylogeny of mammalian host species so that the three major protoparvovirus lineages emerge in concert with major groups of mammalian hosts (**Fig. 5c**). Moreover, we propose a parsimonious model of protoparvovirus evolution wherein: (i) ancestral protoparvovirus species were present in Pangaea prior to its breakup; (ii) the vicariance-driven, deep divergences in the mammalian phylogeny drove the emergence of distinct protoparvovirus lineages in distinct biogeographic regions throughout the course of the Cenozoic Era (from 65 Mya to present); (iii) founding populations of New World rodents were exposed to infection with ApPVs following rodent colonisation of the South American continent (estimated to have occurred ∼50-30 Mya [31]). We found that ancestral range reconstruction, in which a time-calibrated phylogeny of protoparvovirus host species and the present continental distribution of host organisms was used to model the ancestral biogeographical range of protoparvovirus host species [33], supported these findings (**Fig S10a**).

### Recombination and inter-order transfer among Neoprotoparvoviruses

Phylogenies reveal two robustly supported sub-clades within the “Neoproto” clade, one contains the ‘classical’ protoparvoviruses isolated in the 1960s such as Kilham rat virus (KRV) and carnivore protoparvovirus [2], while the other contains more recently identified viruses [34] (**Fig. S7h, S8c**). Three EPVs closely related to these viruses have been reported previously in rodents [14, 35]. Here, a fourth such EPV comprising a near complete parvovirus genome was identified in the steppe mouse (*Mus spicelagus*) (**Fig. 2**). Notably, the NS and VP genes of this element exhibit distinct phylogenetic relationships – in phylogenies based on the *rep* gene (**Fig. S8c**) *proto*.*4-MusSpi* clade groups basal to a clade comprised exclusively of rodent viruses except for one bat-associated virus (BtHp-PV) [36] that emerges as a derived taxon. In contrast, phylogenies based on the VP/Cap gene group BtHp-PV and *proto*.*4-MusSpi* as closely related sister taxa (**Fig. S8c, Fig S10b)**. This implies a history of recombination among rodent parvoviruses in which swapping of VP/Cap genes among lineages has enabled zoonotic transfer into other mammalian groups. It is also notable that protoparvovirus-derived EPVs found in rodent genomes group with viruses isolated from carnivores, bats, and ungulates, rather than those isolated from rodents, and that rodent-associated taxa are interspersed throughout the “Neoproto” clade. This suggests that zoonotic transfer from rodents to other mammalian orders has likely been a defining feature of their evolution.

### Ancient origins of the amdoparvoviruses

We previously reported amdoparvovirus-derived EPVs [32]. Here, we identified novel orthologous copies of an amdoparvovirus-related EPV previously reported in the pit viper (*Protobothrops mucrosquamatus*), demonstrating that it originated >100 Mya (**Table 3, Table S6**). Whereas previous studies had suggested that *EPV-Amdo*.*101-Serpentes* might represent an intermediate lineage between the *Amdo-* and *Protoparvovirus* genera, the Rep phylogenies constructed here placed it robustly with the *Amdoparvovirus* genus. This, combined with the presence of characteristic amdoparvoviral features – e.g., a putative M-ORF and the absence of the PLA2 domain [32] – support the proposed grouping of this EPV within the *Amdoparvovirus* genus and thereby calibrate its evolutionary timeline (**Fig. 4b**). So far, the *Amdoparvovirus* and *Protoparvovirus* genera have only been reported in mammals, suggesting that both groups might have originated in this class, perhaps even relatively recently in (e.g., within the past 20 My). However, our discoveries of: (i) a basal, protoparvovirus-like sequences in a ray-finned fish (**Fig. S9d**) and (ii) a basal and extremely ancient amdoparvovirus-derived EPV (*Amdo*.*101-Serpentes*) in a squamate reptile (**Fig. S3b**) argue for a more distant evolutionary separation between these two genera (**Table 3**).

### Inter-class transmission and the evolution of dependoparvoviruses

Viruses in genus *Dependoparvovirus* can be separated into two distinct groups – one containing the non-autonomous dependoparvoviruses – often referred to as “adeno-associated viruses” (AAVs) - and another containing the autonomous dependoparvoviruses of birds and reptiles [24, 37]. The AAVs are not associated with any disease and are characterised by the requirement for a helper virus to replicate - typically a nuclear DNA virus (e.g., herpesvirus, adenovirus [1]). Dependoparvovirus species lacking this characteristic (‘autonomous parvoviruses’) have been identified in birds and reptiles and are associated with a variety of disease syndromes in ducks and geese [24].

We identified 213 dependoparvovirus-related EPV sequences in mammals, which we estimate to represent at least 80 distinct germline incorporation events (**Table S7**). We reconstructed the evolutionary relationships between dependo-related EPVs and contemporary dependoparvoviruses (**Fig 5, Fig. S8b, Fig. S11**). As expected, phylogenies that included shorter and more degraded EPVs disclosed relatively low bootstrap support for internal branching relationships (**Fig. S12**). However, phylogenies excluding these sequences revealed several robustly supported subclades within the *Dependoparvovirus* genus (**Fig. 6a**). These included clades exclusive to reptilian species (Sauria-), Australian marsupials (Oceania-), and Boreoeutherian mammals (Neo-). A fourth clade, which we named “Shirdal”, contains DPV taxa derived from both avian and mammalian hosts (**Fig. 6a**). Both the composition of this clade in terms of hosts, and its phylogenetic position relative to other DPV groups, implies a role for interclass transmission between mammals and birds in DPV evolution (**Fig. 6b**). Firstly, in both midpoint-rooted phylogenies, and in phylogenies rooted on the saurian dependoparvoviruses (as proposed by Penzes *et al* [38]), the ‘Shirdal’ clade falls intermediate between two exclusively mammalian groups – the non-autonomous AAVs found in placental mammals, and clade Oceania-found exclusively in Australian marsupials (**Fig. 6a**). This implies an ancestral switch from mammalian to avian hosts (green arrow; **Fig. 6b**). Furthermore, the avian viruses in this clade group basally (Ave-), forming a paraphyletic group relative to a derived subclade (Lemuria-) of EPVs obtained from a diverse range of mammalian hosts. This implies a second, subsequent jump from birds to mammals (blue arrow **Fig. 6b**).

**Figure 6.**
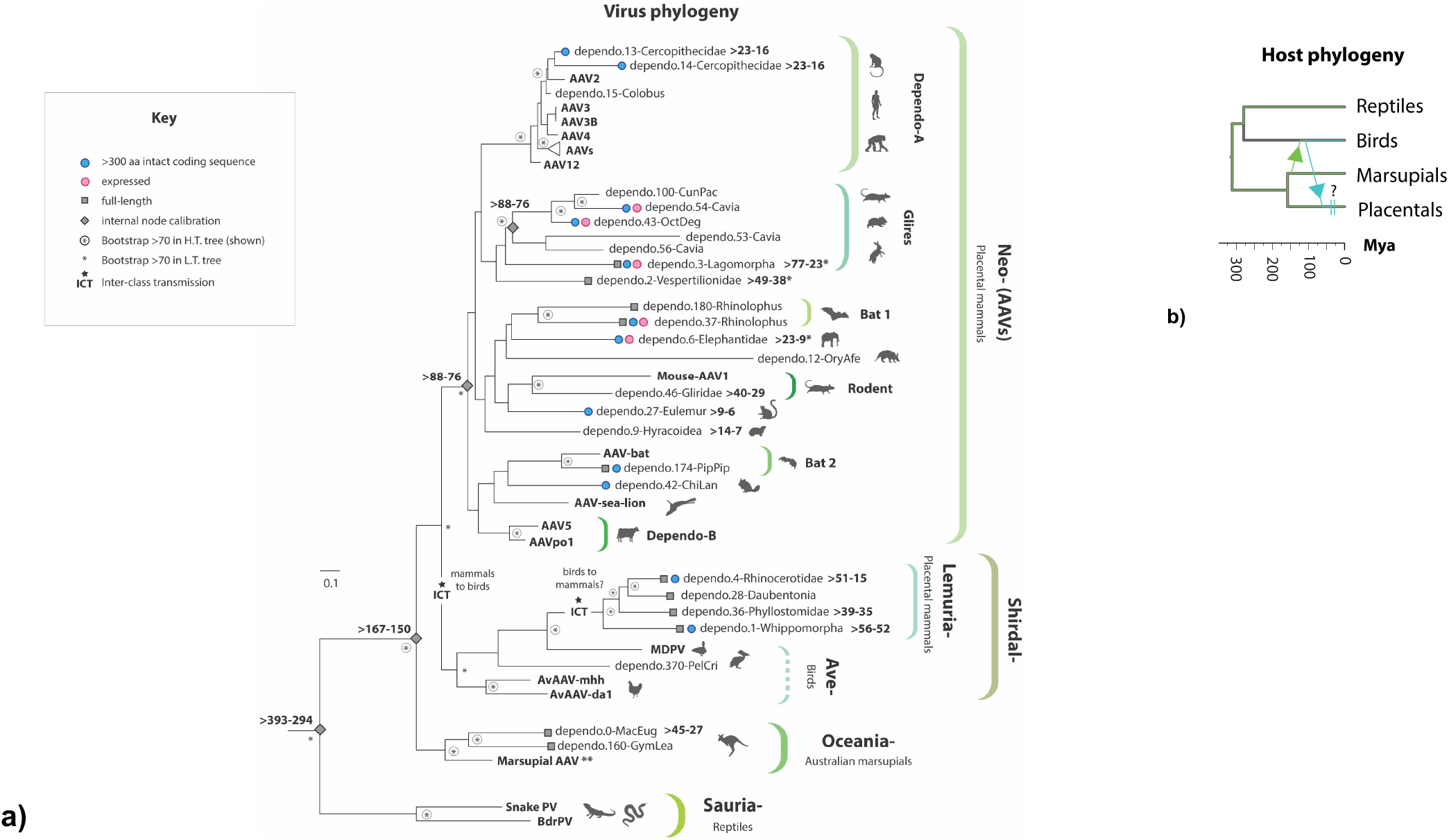
Dependoparvovirus evolution and the influence of inter-class transmission. (a) A maximum likelihood phylogeny showing the reconstructed evolutionary relationships between contemporary dependoparvovirus species and the ancient dependoparvovirus species represented by endogenous parvoviral elements (EPVs). Virus taxa names are shown in bold, EPVs are shown in regular text. The phylogeny was constructed from a multiple sequence alignment spanning (MSA) 330 amino acid residues of the Rep protein and the LG likelihood substitution model and is rooted on the reptilian lineage. Brackets to the right indicate proposed taxonomic groupings. Shapes on leaf nodes indicate full-length EPVs and EPVs containing intact/expressed genes (see key). Numbers next to leaf nodes indicate minimum age calibrations for EPV orthologs. Shapes on branches and internal nodes indicate different kinds of minimum age estimates for parvovirus lineages, as shown in the key. Numbers adjacent to node shapes show minimum age estimates in millions of years before present. For taxa that are not associated with mammals, organism silhouettes indicate species associations, following the key. The scale bar shows evolutionary distance in substitutions per site. Asterisks in circles indicate nodes with bootstrap support >70% (1000 replicates) in the tree shown. Plain asterisks next to internal nodes indicate nodes that are not supported in the tree shown here but do have bootstrap support >70% (1000 replicates) in phylogenies based on longer MSA partitions within Rep (but including less taxa). *Age calibrations based on data obtained in separated publications – see references [18] and [41]. **A contemporary virus derived from the marsupial clade has been reported in marsupials, but only transcriptome-based evidence is available [17]. (b) A time-calibrated phylogeny of vertebrate lineages showing proposed patterns of inter-class transmission in the Shirdal-lineage. **Abbreviations**H.T. tree=high taxa number tree based on shorter alignment (190aa) with more taxa. L.T. tree=low taxa number tree based on longer alignment (350aa). PV=Parvovirus; AAV=Adeno-associated virus; BrdPV=Bearded dragon parvovirus; MdPV=Muscovy duck parvovirus; ORF=open reading frame; aa=amino acid residues.

The ‘Neodependoparvovirus’ clade includes all AAVs, and the EPV fossil record supports the view that the host range of this group encompasses all placental mammals. In Rep phylogenies the AAVs emerge as a derived clade with the autonomously replicating avian viruses grouping basal (**Fig. 6a**). All viruses in this clade fall within the range of diversity defined by two AAV groups (Dependo-A and Dependo-B) both of which are known to exhibit the requirement for a helper virus. This implies that dependency is an ancestral, shared characteristic of AAVs.

### Ancient origins and inter-order transmission of erythroparvoviruses

We identified the first reported examples of EPVs derived from genus *Erythroparvovirus* in the genomes of the Patagonian mara (*Dolichotis patagonum*) - a New World rodent – and the Indri (*Indri indri*), a Malagasy primate (**Fig. 2**). Both EPVs grouped with erythroparvoviruses derived from rodents in phylogenetic trees, indicating inter-order transmission from rodents to lemuriforme primates (**Fig. S13**). Furthermore, when examined in relation to the biogeographic distribution of host species, these phylogenetic relationships provide tentative age calibrations for the *Erythroparvovirus* genus based on the parsimonious assumption that they spread to Madagascar and South America during the Cenozoic Era along with rodent founder populations (**Table 3**).

### Ancient origins of vertebrate hamaparvoviruses (subfamily Hamaparvovirinae)

The recently defined subfamily *Hamaparvovirinae* contains two genera known to infect vertebrates – *Chaphamaparvovirus* and *Ichthamaparvovirus* [1, 39]. Our previous studies have shown that chaphamaparvovirus-derived sequences occur in WGS data, but all seem to derive from contaminating viruses rather than EPVs [22, 39] (**Table S11**). However, we previously identified *Ichthamaparvovirus*-derived EPVs in fish [39], and here report an additional *Ichthamaparvovirus*-derived EPV locus in snakes (suborder Serpentes) (**Table S10**). Importantly, this sequence demonstrates that the host range of this virus genus extends to reptiles (**Fig 2**., **Fig. S14**). Furthermore, we show that orthologs are present in multiple snake species, establishing a minimum age of 62 My for the *Ichthamaparvovirus* genus (**Table 3**).

### Coding capacity and expression of EPV sequences

Previous studies have shown that some EPV loci express RNA with the potential to encode polypeptide gene products, either as unspliced viral RNA [17, 40, 41], or as fusion genes comprising RNA sequences derived from both host and viral sources [42]. We examined all EPV loci identified in our study to determine their coding potential. EPV loci ranged from near full-length virus genomes to short fragments ∼300 nucleotides (nt) in length (**Fig. 2**). We identified 25 unique loci at which sequences encoding uninterrupted polypeptide sequences of 300 amino acids (aa) or more are retained (**Table 4**). Long stretches of intact coding sequence have been maintained in a diverse collection of EPVs, including some that are demonstrably many millions of years old (e.g., *Amdo*.*101-Serpentes*). Notably, we identified EPVs in cercopithecine primate genomes that appear to represent the ancient progenitors of contemporary primate AAVs, including one (*Dependo*.*14-Cercopithecidae*) that retains 309 aa of NS/Rep despite originating ∼23-16 Mya (**Table 4, Table S7**). For several intact EPVs >300 aa in length, mRNA expression has been experimentally demonstrated [18, 40, 41] while screening of RNA databases revealed evidence for expression of EPV RNA in several additional EPVs (**Table 4**).

**Table 4.** Coding capacity and expression of vertebrate EPVs.

| Genus | Clade<sup>a</sup> | Number<sup>b</sup> | Organism<sup>c</sup> | Exp.<sup>d</sup> | Length<br>(aa)<sup>e</sup> | Genes<sup>f</sup> |
| --- | --- | --- | --- | --- | --- | --- |
| Amdo- |  | 1 | *Ellobius lutescens* | *p* | 383 | NS |
| Amdo- |  | 2 | " |  | **586** | NS-MAFG* |
| Amdo- |  | 101 | *Protobothrops mucrosquamatus* | *p* | **574** | NS |
| Amdo- |  | 101 | " |  | 333 | **VP** |
| Dependo- | Oceania- | 0 | *Macropus eugenii* |  | 442 | **VP** |
| Dependo- | Lemuria- | 1 | *Orcinus orca* |  | 357 | **VP** |
| Dependo- | Lemuria- | 1 | *Globicephala melas* |  | 314 | NS |
| Dependo- | Neo- (AAV) | 2 | *Myotis lucifugus* | *p* | 311 | NS |
| Dependo- | Neo- (AAV) | 3 | *Lepus timidus* | *c* | 439 | NS |
| Dependo- | Neo- (AAV) | 4 | *Dicerorhinus sumatrensis* |  | 321 | **VP** |
| Dependo- | Neo- (AAV) | 6 | *Loxodonta africana* | *c* | **583** | NS |
| Dependo- | Neo- (AAV) | 13 | *Cercocebus atys* |  | 309 | NS |
| Dependo- | Neo- (AAV) | 14 | " |  | **561** | NS |
| Dependo- | Neo- (AAV) | 27 | *Eulemur macaco* |  | 473 | NS |
| Dependo- | Neo- (AAV) | 37 | *Rhinolophus sinicus* |  | 382 | NS |
| Dependo- | Neo- (AAV) | 42 | *Chinchilla lanigera* |  | **499** | NS |
| Dependo- | Neo- (AAV) | 43 | *Octodon degus* | *c* | **502** | NS |
| Dependo- | Neo- (AAV) | 54 | *Cavia porcellus* | *c* | **230** | NS-Myo9* |
| Dependo- | Neo- (AAV) | 59 | *Myocastor coypus* |  | 304 | NS |
| Dependo- | Neo- (AAV) | 88 | *Megaderma lyra* |  | 330 | NS |
| Dependo- | Neo- (AAV) | 174 | *Pipistrellus pipistrellus* |  | 449 | NS |
| Dependo- | Neo- (AAV) | 174 | " |  | **634** | **VP** |
| Dependo- | Sauria- | 201 | *Thamnophis elegans* |  | **722** | **VP** |
| Dependo- | Sauria- | 201 | " |  | 385 | **VP** |
| Dependo- | Sauria- | 202 | " |  | 404 | **VP** |
| Erythro- |  | 1 | *Indri indri* |  | **701** | NS |
| Erythro- |  | 1 | " |  | **567** | **VP** |
| Ichthama- |  | 2 | *Emydocephalus ijimae* |  | 478 | ORF2 |
| Ichthama- |  | 2 | *Hydrophis melanocephalus* |  | 392 | NS |
| Proto- | Neo- | 1 | *Rattus norvegicus* |  | 440 | **VP** |
| Proto- | Neo- | 2 | *Mus spretus* |  | 301 | NS |
| Proto- | Neo- | 2 | " |  | 484 | **VP** |
| Proto- | Neo- | 3 | *Apodemus sylvaticus* |  | 341 | **VP** |
| Proto- | Neo- | 3 | " |  | 314 | NS |
| Proto- | Neo- | 4 | *Mus spicilegus* |  | 403 | NS |
| Proto- | Neo- | 4 | " |  | **714** | **VP** |
| Proto- | Meso- | 102 | *Tamandua tetradactyla* |  | **563** | NS |
| Proto- | Meso- | 102 | " |  | **686** | **VP** |
| Proto- | Archaeo- | 103 | *Monodelphis domestica* |  | 349 | **VP** |
| Proto- | Archaeo- | 107 | *Myocastor coypus* |  | 486 | NS |
| Proto- | Archaeo- | 107 | " |  | **589** | **VP** |
| Proto- | Archaeo- | 108 | *Hydrochoerus hydrochaeris* |  | 371 | NS |
| Proto- | Archaeo- | 131 | *Ctenomys sociabilis* |  | 346 | NS |
| Proto- | Archaeo- | 131 | " |  | 473 | **VP** |
| Proto- | Archaeo- | 134 | *Dolichotis patagonum* |  | 337 | NS/VP |
| Proto- | Archaeo- | 134 | " |  | 387 | **VP** |
| Proto- | Archaeo- | 138 | *Hydrochoerus hydrochaeris* |  | 429 | **VP** |
| Proto- | Archaeo- | 139 | *Myocastor coypus* |  | 334 | **VP** |
| Proto- | Archaeo- | 141 | *Erethizon dorsatum* |  | 361 | NS |
| Proto- | Archaeo- | 143 | *Cuniculus paca* |  | 462 | NS/VP |
| Proto- | Archaeo- | 152 | *Sarcophilus harrisii* |  | 363 | NS |
| Proto- | Archaeo- | 155 | *Gymnobelideus leadbeateri* |  | 320 | **VP** |
| Proto- | Archaeo- | 158 | *Macropus eugenii* |  | 429 | **VP** |
| Proto- | Archaeo- | 178 | *Erethizon dorsatum* |  | 308 | NS |
| Proto- | Archaeo- | 189 | *Grammomys surdaster* | *p* | **82** | NS |
| Proto- | Archaeo- | 210 | *Macropus eugenii* |  | 327 | **VP** |
| Proto- | Archaeo- | 210 | " |  | 314 | NS |

of coding sequence or have evidence of expression. Sequences that are <300 aa in length but with evidence for expression are underlined. <sup>†</sup> VP shown in bold. \* EPV-gene fusion products predicted [32] or confirmed [42] – length shown is for viral portion only.

## DISCUSSION

The accumulation of genome sequence data from diverse, novel parvovirus species and EPVs presents unprecedented opportunities to investigate parvovirus biology using comparative approaches. Here, we sought both to develop a more detailed view of the long-term co-evolutionary interactions between parvoviruses and vertebrates, and to establish tools and data standards that facilitate the implementation of broad-scale, comparative analyses of parvovirus genomes. We recovered the complete repertoire of EPV sequences in WGS data representing 752 vertebrate species. While previous studies have reported a sampling of EPV diversity in vertebrates [12-18, 22, 32, 35, 40, 42, 43], our investigation is an order of magnitude larger in scale: we identify 364 sequences representing nearly 200 discrete germline incorporation events that took place during the Cenozoic Era (**Fig. 2, Table S1**). Furthermore, we introduced a higher level of order to these data by: (i) discriminating between unique loci and orthologous copies; (ii) aligning EPVs to contemporary viruses and hierarchically arranging MSAs so that phylogenetic and genomic comparisons can utilise the maximum amount of available data; (iii) applying a standardised nomenclature to EPVs that captures information about orthology and taxonomy.

Our analysis shows that parvovirus DNA was incorporated into the vertebrate germline throughout the Cenozoic Era (**Fig. 3**). The independent formation and fixation of EPVs in such a diverse range of vertebrate groups suggests that *Parvovirinae* genera circulated widely among vertebrate fauna throughout their evolution. Furthermore, the robust calibrations we obtain from EPVs lend credibility to similarly ancient, biogeography and distribution-based age estimates obtained for other parvovirus lineages, such as the Ave-Boca lineage, that are not represented in the genomic ‘fossil record’ of parvoviruses (**Fig. 4, Table 3**). Given that the origins of the parvovirus family likely extend far back into the early evolutionary history of animal species [16, 44] we imagine that the subfamily *Parvovirinae*, which includes ten vertebrate-specific genera, could have evolved in broad congruence with the emergence and diversification of major vertebrate groups. Indeed, this extended evolutionary timeline can be strikingly visualised in the *Protoparvovirus* genus, in which the emergence and spread of sub-lineages reflects the impact of vicariance and continental drift on mammal evolution during the Cenozoic Era (**Fig 4**.).

In relation to this, it is notable that the same host species and species groups harbour infections with multiple distinct genera of parvoviruses - for example, at least seven distinct genera circulate in mammals. Furthermore, EPV-based calibrations indicate that these genera are likely to have co-circulated among mammals for many millions of years (**Fig. 3**). Ecology is thought to be an important factor in speciation, because natural selection can drive adaptive divergence between lineages that inhabit different environments [45]. The extended evolutionary timeline implied our analysis supports the idea that the persistence of multiple, distinct parvovirus genera in the same host species groups reflects adaptive divergence among these genera, such that each parvovirus occupies a distinct ‘ecological niche’ (i.e., part of the ecological space available in the environment). Although niches can be difficult to define precisely, most treatments consider conditions of the physical environment, characteristics of resources, and the traits of other interacting species as important factors [46]. In the case of viruses, host species range is inevitably a major influence, but there are likely many other aspects of replication that come into play. While all members of the subfamily *Parvovirinae* use similar basic mechanisms to achieve specific steps in infection, the details of these processes (e.g., the specific cell types and organ systems targeted for replication) frequently differ between genera. For example, primate erythroparvoviruses target erythroid progenitor cells [3], mammalian chaphamaparvoviruses are suspected to be nephrotropic [47], and antibody-dependent enhancement is suspected to a shared characteristic of amdoparvoviruses [48, 49].

The non-autonomous parvoviruses or ‘AAVs’ provide a stand-out example of adaptation to a specialised niche. It may seem counterintuitive to regard their requirement for a helper virus as an adaptation, since it may appear superficially to be a deficiency. However, molecular genetic analysis of AAV2 indicates that helper-virus dependent replication resulted from a gain of function rather than loss of autonomous replication. Simplified, the functional gains were additional levels of control over virus gene expression. For example, the cellular transcription factor YY1 functions as a transcriptional repressor that inhibits expression of the AAV2 p5 promoter and the large Rep proteins, but in the presence of adenovirus co-infection and E1a gene product(s), YY1 becomes a transcriptional activator leading to p5 transcription and large Rep protein expression [50, 51]. Presumably, a similar relationship exists between YY1 and other helper-viruses leading to transcriptional activation of the dependoparvovirus NS proteins. The gain of dependency also attenuates the pathogenicity of the virus: without helper virus co-infection AAV replication is suppressed, and cells appear unaffected by the presence of virus DNA.

Conceivably, dependency could have evolved as a means of timing replication to optimise the probability of successful transmission. The double-stranded DNA (dsDNA) virus groups that facilitate AAV replication (e.g., adenoviruses, herpesviruses) include species that can ‘sense’ the cellular environment and switch to replicative mode when conditions are optimal [52]. By ‘tethering’ their replication to that of larger and more sophisticated dsDNA viruses AAVs can potentially take advantage of the ability of these viruses to: (i) optimise the timing of replication, and (ii) establish a cellular environment favourable to virus replication. In relation to this, it is intriguing that one betaherpesvirus genus (*Roseolovirus*) has acquired a homolog of the parvovirus Rep gene (U94) that can reportedly rescue replication of AAV [53], particularly since - like dependoparvoviruses - roseoloviruses can establish latent infections by integrating their genomes into chromosomal DNA of infected cells [54]. U94 is dispensable for integration, but the retention of this gene in multiple species (**Fig. S9**) indicates that it plays a yet undetermined role in the betaherpesvirus life cycle.

While inter-class transmission of parvoviruses appears to be rare overall, we obtained compelling evidence that it has occurred relatively frequently among autonomous dependoparvoviruses (**Fig. 6**). Phylogenies imply that zoonotic transfer of these viruses from marsupials to birds, then subsequently from birds into placental mammals. Both of these inter-class transmission events must have occurred in the distant evolutionary past, as they pre-date the emergence of spread of the ‘Lemuria’ clade – a group that is represented only by EPV ‘fossils’ and includes some of the oldest EPV loci yet identified (**Fig. 4, Table 3**) [18]. Interestingly, recent studies have reported the spread of autonomous parvoviruses related to MDPV and GPV from anseriform birds to the relatively distantly related ratites (flightless birds) [55], suggesting that these viruses might have an inherent capacity to cross between relatively distantly related host groups, and may represent emerging pathogens of poultry and wild birds.

The extent to which vertebrate EPVs have reached fixation through positive selection as opposed to incidental factors such as founder effects, population bottlenecks, and genetic hitchhiking, remains unclear. Potentially, EPVs might sometimes be co-opted or “exapted” as has been reported for EVEs derived from other virus groups, including retroviruses (family *Retroviridae*) [56-58], bornaviruses (family *Bornaviridae*) [59], and polydnaviruses (family *Polydnaviridae*) [60]. Recent studies have revealed that two distinct, fixed EPVs in the germline of the degu (*Octodon degus*) and (ii) family Elephantidae (elephants) encode an intact Rep protein ORF and also exhibit similar patterns of tissue-specific expression in the liver [40, 41], suggesting that expression of Rep protein or mRNA from EPV loci may be physiologically relevant. Experimental studies have shown that AAV Rep protein (over) expression affects healthy cells through a variety of activities including DNA binding, constitutive ATPase, and inhibiting the (cyclic) cAMP-activated protein kinase A (PKA) and protein kinase X (PrKX) [61, 62]. Rep-mediated inhibition of these kinases not only affects the infected cell, but also diminishes the proliferation of adenovirus helper virus, perhaps attenuating virus-induced pathogenesis [63, 64]. Conceivably, some of these properties that have favoured capture of Rep genes in host species and betaherpesvirus genomes. EPVs are also commonly derived from the VP/capsid ORF (**Table S12**), and while it remains unknown if these represent co-opted or exapted sequences, their conservation is quite striking. ORFs derived from the capsid proteins of other virus groups have also been retained in animal genomes and experimental evidence supports the idea that these EVEs can function as antiviral immune factors capable of blocking infection with related viruses [57].

Our investigation shows that vertebrate parvoviruses have ancient associations with their hosts and have acquired lineage-specific adaptations over many millions of years. Importantly, this implies that the growing wealth of genomic data - incorporating both virus and EPV sequences - can be harnessed to define the biological characteristics of parvovirus species and groups. By linking our growing knowledge of parvovirus distribution, diversity, and evolution with experimental and field studies, we can now develop parvovirus-based therapeutics that are grounded in an understanding of natural history. Such comparative, evolutionary approaches will not only help identify virus species with desirable characteristics – e.g., a well-defined tropism for a specific cell type – they establish a powerful, rational basis for dissecting structure- function relationships in parvovirus genomes. Evolution provides a unifying framework that naturally links knowledge discovery efforts, regardless of the biological scale(s) being addressed or the analytical context. To support the broader use of evolution-related domain knowledge in parvovirus research we published our data in the form of an open, extensible, database framework (**Fig. S1**). We hope that by enabling the reproduction of comparative genomic analyses and supporting re-use of the complex datasets that underpin them, these resources can help researchers exploit genomic data to develop a deeper understanding of parvovirus biology.

## METHODS

### Creation of resources for reproducible comparative analysis of parvovirus genomes

We used GLUE (Genes Linked by Underlying Evolution), a bioinformatics software framework, to develop open data resources for parvoviruses [65]. The advantages of GLUE include: (i) the organization of virus sequence data in relation to their hypothesized evolutionary relationships (an approach that is key to the practical interpretation of sequences); (ii) integration with wider computer systems, using standard technologies such as MySQL and JSON; (iii) mechanisms to customize functionality on a usage-specific basis, for example schema extension and scripting mechanisms; (iv) features and functionality to streamline rapid development of bespoke analysis projects.

A library of parvovirus reference sequences (**Table S1**) was compiled. We obtained reference genome sequences for all Parvovirus species recognised by the International Committee for Taxonomy of Viruses (ICTV). This set was supplemented with genome sequences representing recently reported parvovirus species that have not yet been incorporated into official taxonomy, and parvovirus-derived EPVs – these were identified via searches of public databases (PubMed, NCBI nucleotide). A standard set of genome features was defined (**Table S2**). We then selected well-annotated reference genomes as ‘master’ reference sequences for each subfamily, genus, and species represented in our MSA hierarchy (see below & **Table 1**) and mapped the location of genome features in master reference genomes. Multiple sequence alignments (MSAs) were constructed using MUSCLE [66] and GLUE’s native BLAST-based aligners [21, 67]. The Parvovirus-GLUE project is built by using GLUE’s native command layer to create a bespoke MySQL database that not only contains the data items associated with our analysis, but also maps the semantic links between them (e.g., the associations between specific sequences, genome features, and MSA segments) (**Fig. S1, Fig. S2**).

Isolate data was captured by extending GLUE’s underlying database schema. GenBank sequences in XML format are imported into the Parvovirus-GLUE project using GLUE’s ‘genbankImporter’ module to extract sequence and isolate-associated data. Non-standard fields (e.g., isolate-specific information in the ‘notes’ section of the GenBank entry) were extracted using a regular expression library, and a FreeMarker template was used to standardise their values as described previously [19]. Additional data (e.g., information missing from GenBank records but identified in an associated publication) was imported from tabular files using GLUE’s TextFilePopulator module. Where (non-master) reference genome sequences were lacking feature annotations, we used GLUE’s ‘inherit features’ command [19] to infer their coordinates from an MSA in which the genomic coordinate space was constrained to a fully annotated master reference sequence for the corresponding genus (see **Table 1**).

When performing comparative sequence analyses it is often practical to construct separate MSAs representing distinct taxonomic levels (e.g., family, subfamily, genus). However, maintaining such MSA ‘ensembles’ so that they remain cohesive with one another (i.e., such that the implied homologous relationships are consistent across all alignments that share two or more members) is a labour-intensive task. To address this, GLUE allows MSAs to be hierarchically linked via a ‘constrained alignment tree’ data structure [19] (**Table 1, Fig. S3**). In each MSA a chosen master reference sequence constrains the genomic coordinate space. For contemporary parvoviruses we used our chosen genus master reference sequences, while for EPVs we used consensus EPVs as constraining references for MSAs representing orthologous EPV loci. Each MSA in the Parvovirus-GLUE MSA hierarchy is linked to its child and/or parent MSAs via our chosen set of references (**Table 1**) – this creates in effect a single, unified MSA that can be used to implement comparisons across a range of taxonomic levels while also making use of the maximum amount of available information at each level. By standardising the genomic coordinate space to the constraining reference sequence selected for each MSA, the alignment tree enables standardised genome comparisons across the entire *Parvoviridae* family. Note that, while MSAs representing internal nodes (see **Table 1**) contain only master reference sequences, they can be recursively populated with all taxa contained in child alignments when exported from GLUE for analysis [19].

To accommodate EPV data in Parvovirus-GLUE, we extended the underlying database schema to incorporate an EPV-specific table with data fields capturing EPV characteristics (e.g., locus coordinates, ortholog group, flanking genes). In addition, where multiple EPV orthologs were identified, we created MSAs to represent homology between individual orthologs in each EPV set, we used these to: (i) reconstruct the evolutionary relationships between orthologs [20];

(ii) derive consensus reference sequences for each EPV locus.

We used Parvovirus-GLUE to implement a process for reconstructing evolutionary relationships across the entire *Parvoviridae* - first among viruses (**Fig. S6**) and secondly among viruses and EPVs (**Fig. S6**). MSAs partitions derived from the constrained MSA tree (**Table 1**) were used as input for phylogenetic reconstructions. To handle coverage-related issues we used GLUE’s ‘record feature coverage’ functionality to generate gene coverage data for all aligned sequences prior to phylogenetic analysis and then used this information to condition the way in which taxa are selected into MSA partitions.

The GLUE software framework broadly adheres to a ‘data-oriented programming’ paradigm in which an explicit separation is maintained between code and data, and data is treated as immutable – this approach directly addresses issues of reusability, complexity, and scale in the design of information systems [68]. To illustrate how Parvovirus-GLUE can be used to perform standardised, reproducible, comparative genomic analyses comparative analysis of parvovirus genomes, we implemented a simple script that uses information collated in Parvovirus-GLUE to derive nucleotide, dinucleotide, and amino acid composition metrics for all parvovirus species, broken down by genome feature (**Table S4, Fig. S4**).

Parvovirus-GLUE can be used as a local, stand-alone tool or as a foundation for the development of genome analysis-based reporting tools for potential use in human and animal health - e.g., see HCV-GLUE [19], RABV-GLUE [69]. In the interests of maintaining a lightweight, approach, the published project contains only a single reference genome for each parvovirus species. However, it can readily be expanded to focus on analysis at species level. A tutorial included with the published resource, demonstrates how this can be done [20]. The Parvovirus-GLUE project can be installed on all commonly-used computing platforms, and is also fully containerised via Docker [70]. Finally, hosting of the Parvovirus-GLUE resource in an openly accessible online version control system (GitHub) provides a platform for managing its ongoing development, following practices established in the software industry (**Fig. S1c**) [71]. We hope that these resources will enable researchers working in different areas of parvovirus genomics to benefit from one another’s work.

### in silico screening of whole genome sequence databases

We used the Database-Integrated Genome Screening (DIGS) tool [72] to derive a non-redundant database of EPV loci within published WGS assemblies. The DIGS tool is a Perl-based framework in which the Basic Local Alignment Search Tool (BLAST) program suite [67] is used to perform similarity searches and the MySQL relational database management system to coordinate screening and record output data. A user-defined reference sequence library provides (i) a source of ‘probes’ for searching WGS data using the tBLASTn program, and (ii) a means of classifying DNA sequences recovered via screening (**Fig. S4)**. For the purposes of the present project, we collated a reference library composed of polypeptide sequences derived from representative parvovirus species and previously characterised EPVs. WGS data of animal species were obtained from the National Center for Biotechnology Information (NCBI) genome database [73]. We obtained all animal genomes available as of March 2020. We extended the core schema of the screening database to incorporate additional tables representing the taxonomic classifications of viruses, EPVs and host species included in our study. This allowed us to interrogate the database by filtering sequences based on properties such as similarity to reference sequences, taxonomy of the closest related reference sequence, and taxonomic distribution of related sequences across hosts. Using this approach, we categorised sequences into: (i) putatively novel EPV elements; (ii) orthologs of previously characterised EPVs (e.g., copies containing large indels); (iii) non-viral sequences that cross-matched to parvovirus probes (e.g., retrotransposons). Sequences that did not match to previously reported EPVs were further investigated by incorporating them into genus-level, genome-length MSAs (see **Table 1**) with representative parvovirus genomes and reconstructing maximum likelihood phylogenies using RAxML (version 8.2.12) [74].

Where phylogenetic analysis supported the existence of a novel EPV insertion, we also attempted to: (i) determine its genomic location relative to annotated genes in reference genomes; and (ii) identify and align EPV-host genome junctions and pre-integration insertion sites. Where these investigations revealed new information (e.g., by confirming the presence of a previously uncharacterised EPV insertion) we updated our reference library accordingly. This in turn allowed us to reclassify EPV loci in our database and group sequences more accurately into categories. By iterating this procedure, we progressively resolved the majority of EPV sequences identified in our screen into groups of orthologous sequences derived from the same initial germline incorporation event (**Table S5-S10**).

### Phylogenetic and Phylogeographic analysis

Nucleotide and protein phylogenies were reconstructed using maximum likelihood (ML) as implemented in RAxML (version 8.2.12) [74]. Protein substitution models were selected via hierarchical maximum likelihood ratio test using the PROTAUTOGAMMA option in RAxML. For multicopy EPV lineages, we constructed MSAs and phylogenetic trees to confirm that branching relationships follow those of host species (**Fig S4b**, [20]). Time-calibrated vertebrate phylogenies were obtained via TimeTree [75].

To model ancestral biogeographical range of protoparvovirus hosts [33], we obtained country-level distribution information for each host species via the occ_search function of the rgbif library in R [76]. Country records were consolidated in continent entries with the continents function of the countrycode library and manually curated to ensure accuracy. Within continents, North Africa and Sub-Saharan Africa were considered distinct distributions and coded separately. The Dispersal-Extinction-Cladogenesis (DEC) model implemented in the program Lagrange C++ was applied without constraining the number of ancestral states nor limiting connectivity between biogeographic units [77]. Ancestral states at all nodes in the tree were inferred and the tree visualized in R with the ggtree and ggplot libraries [78, 79].

### Genomic analysis of EPVs

ORFs were inferred by manual comparison of sequences to those of reference viruses. For phylogenetic analysis, the putative peptide sequences of EVEs (i.e., the virtually translated sequences of EVE ORFs, repaired to remove frameshifting indels) were aligned with polypeptide sequences encoded by reference genomes. We used PAL2NAL [80] to generate in-frame, DNA alignments of virus coding domains from alignments of polypeptide gene products. Phylogenies were reconstructed using maximum likelihood (ML) as implemented in RAxML [74] and GTR model of nucleotide selection as selected using the likelihood ratio test. The putative peptide sequences of EPVs were aligned with NS and VP polypeptides of representative exogenous parvoviruses using MUSCLE (version 3.8.31).

### Expression and intactness of EPVs

We identified open coding regions of coding sequence in EPVs using scripts included with Parvovirus-GLUE [20]. To determine if there was evidence of expression of EPVs in host species, we searched the NCBI Reference RNA Sequences (refseq_rna) with Dependoparvovirus VP and Rep sequences (NC_002077). We searched a translated nucleotide query and a translated database using tBLASTx [67] and evaluated alignments found between refseq_rna sequences and Dependoparvovirus VP and Rep sequences. To further verify expression, we determined if the annotations were solely based on computational prediction or if there is RNAseq data alignment to the annotation in support of the feature. For those host species with evidence of expression, we conducted BLASTn searches within refseq_rna to identify expressed EPVs.

## ACKNOWLEDGEMENTS

This work was supported by funding from the Association Monégasque Contre les Myopathies, and the Bill & Melinda Gates Foundation (OPP1202116). We thank Professor Greg Towers for helpful comments on the manuscript.

