## Supplemental figures for "Comparative analysis reveals the long-term co-evolutionary history of parvoviruses and vertebrates"

**Figure S1.** Parvovirus-GLUE open resource for reproducible comparative analysis of parvovirus genome data.

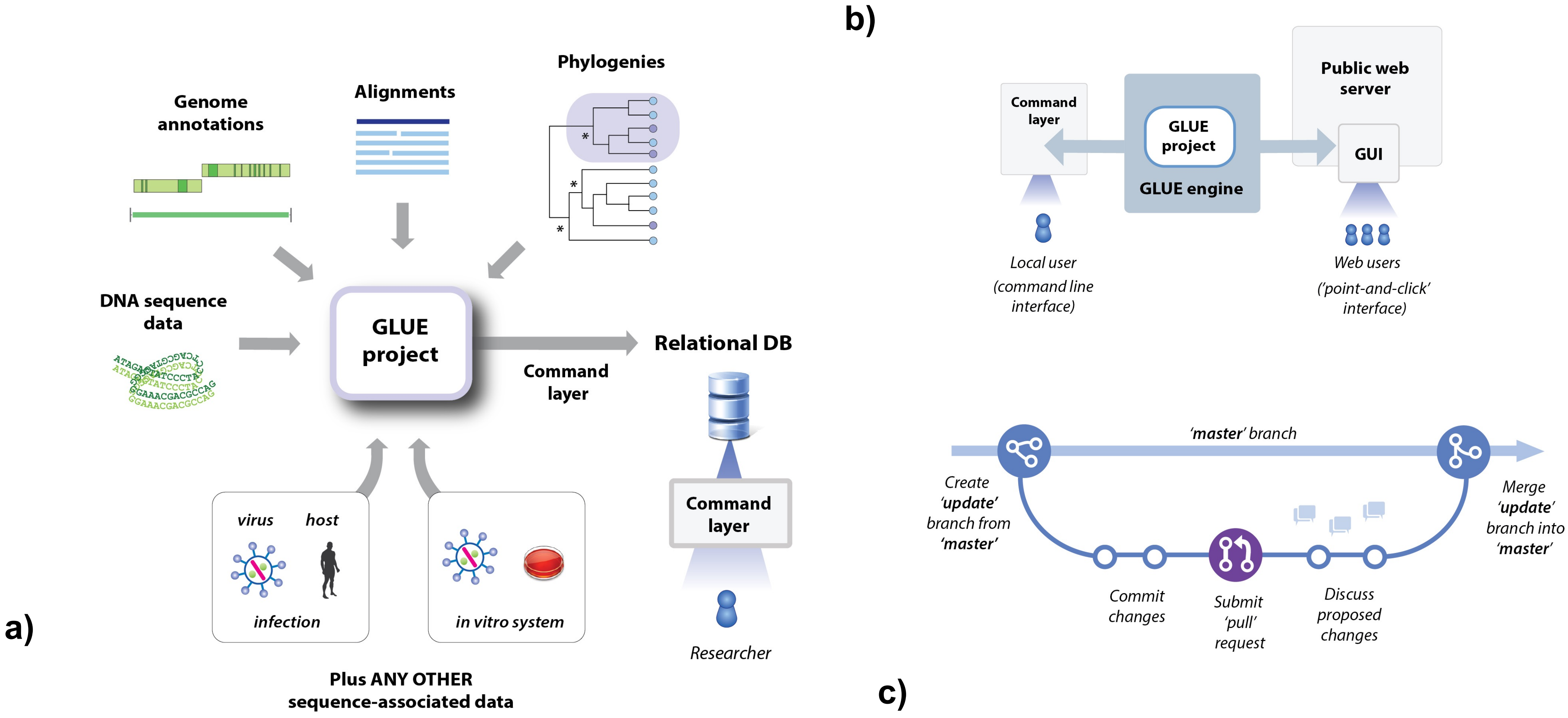

**Figure S2.** The Parvovirus-GLUE resource build process.

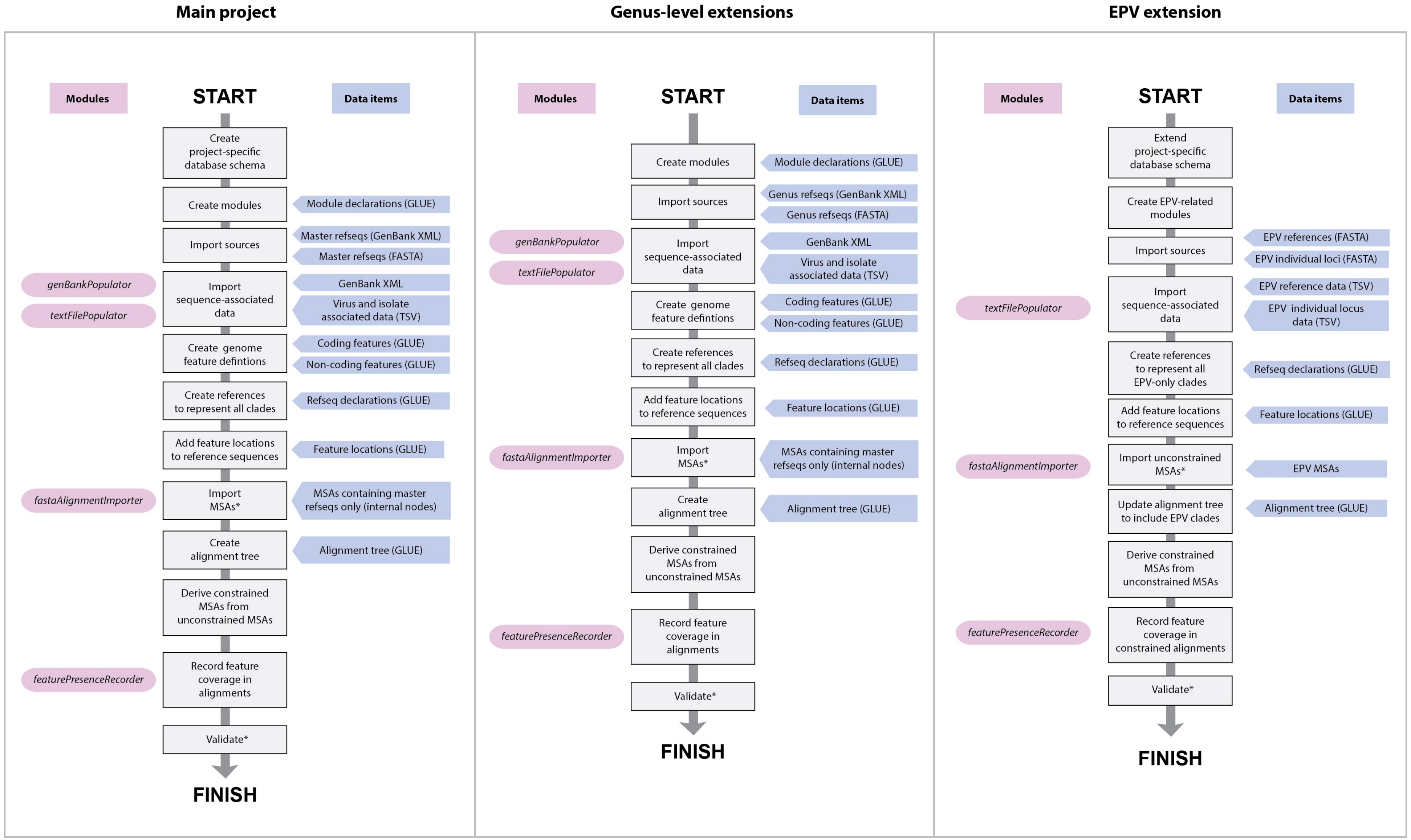

**Figure S3.** The constrained multiple sequence alignment tree in Parvovirus-GLUE.

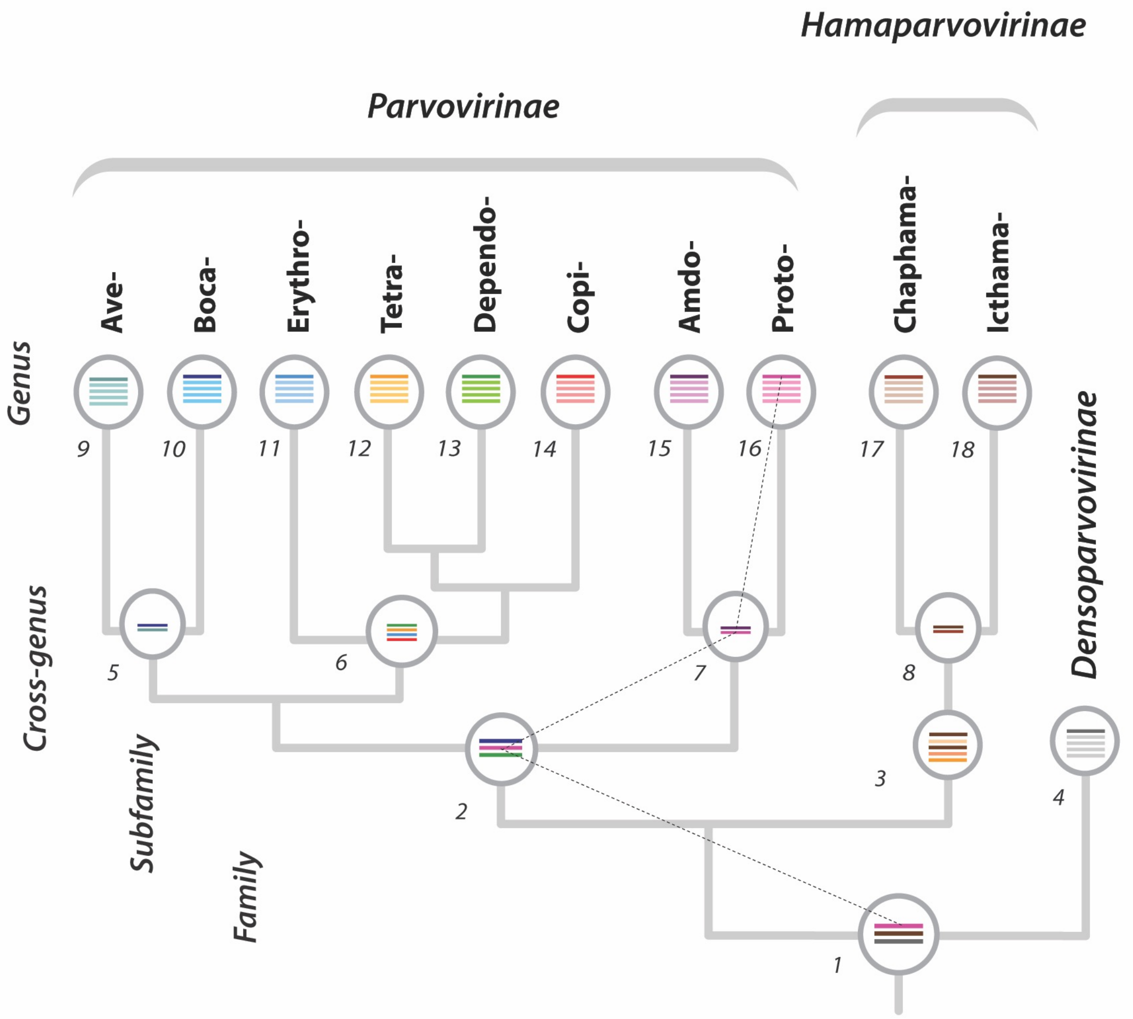

**Figure S4 (a)**

Nucleotide composition

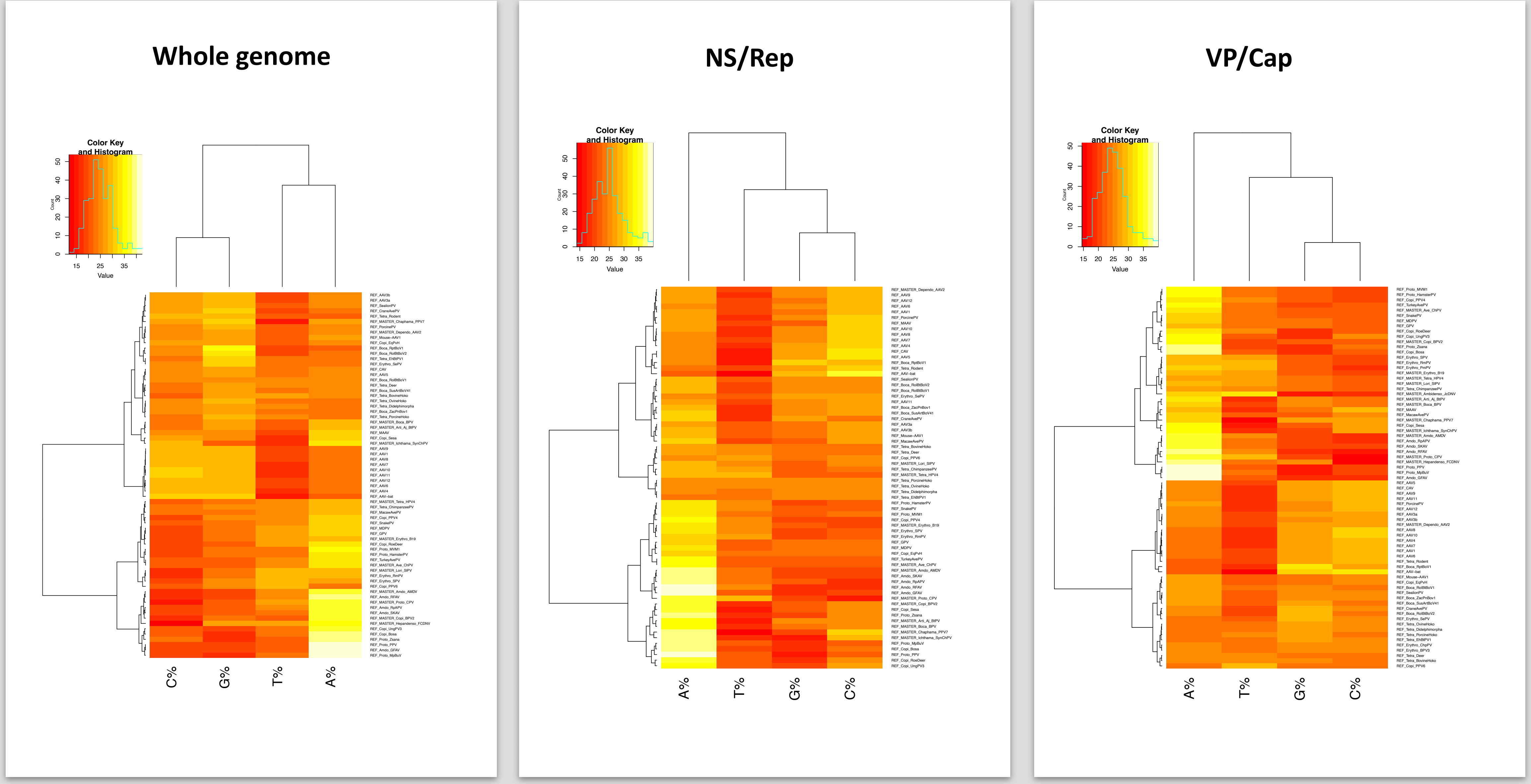

**Figure S4 (b)**

Dinucleotide composition

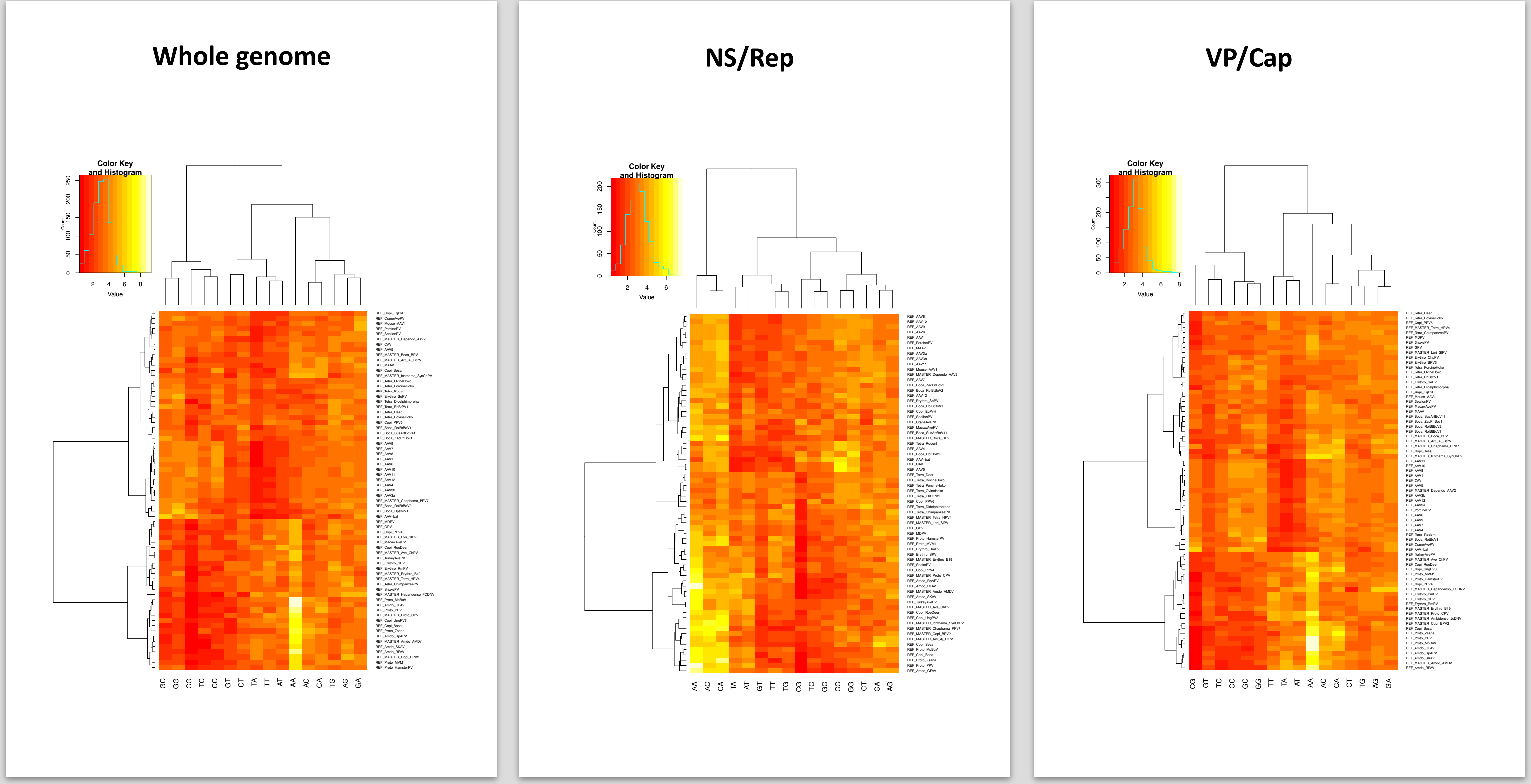

**Figure S4 (c)**

Amino composition

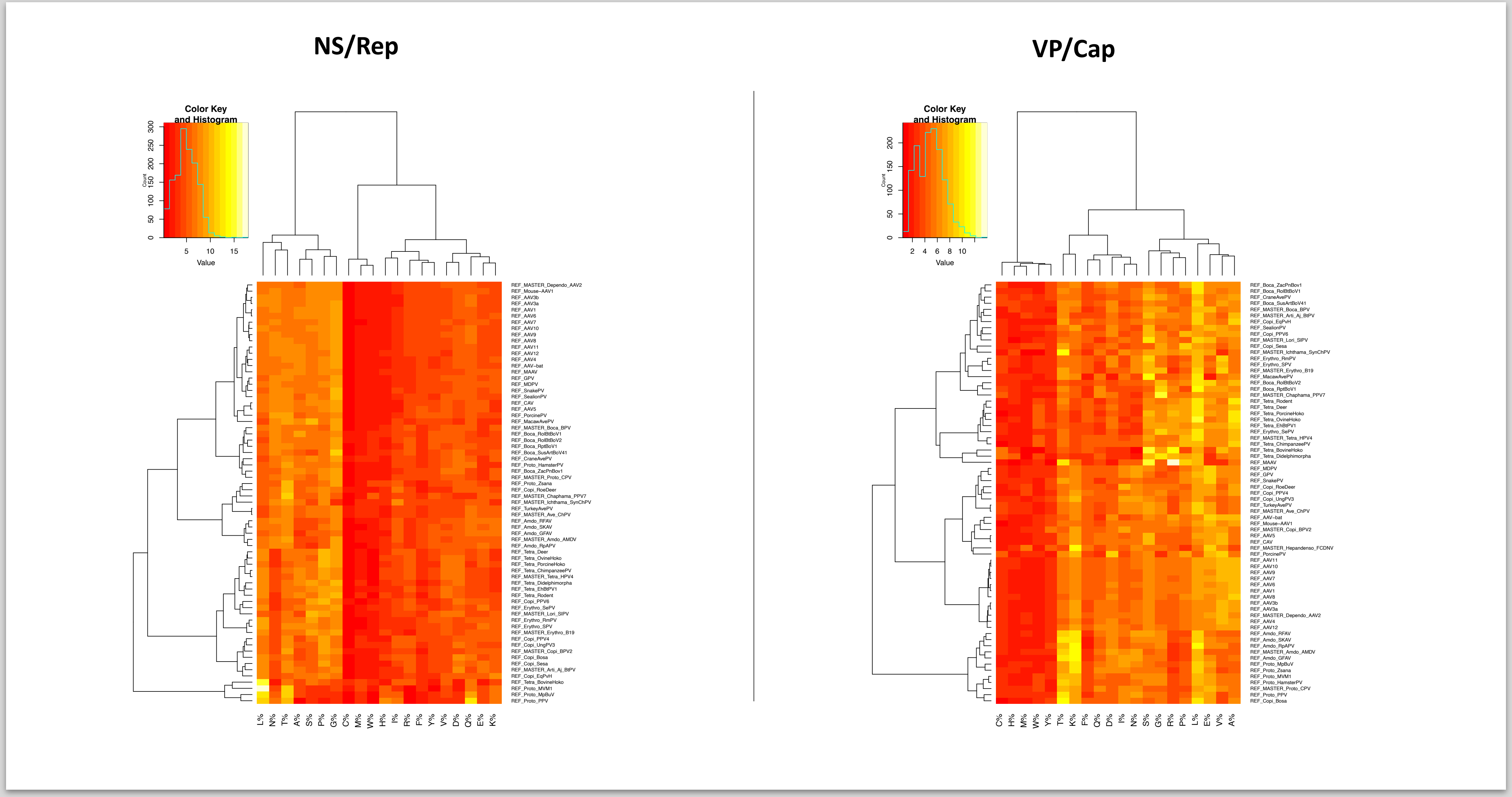

**Figure S5.** Genome screening *in silico.*

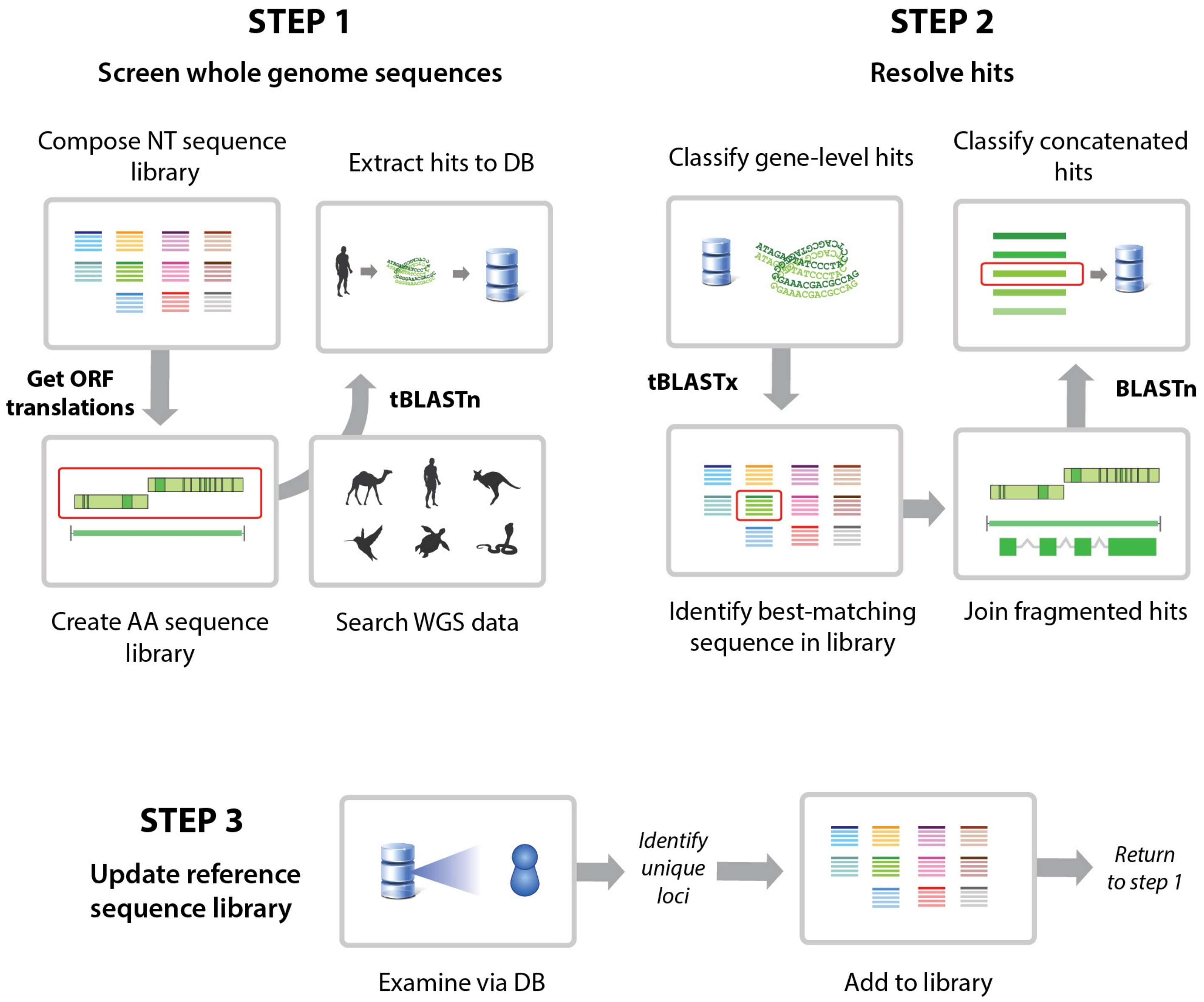

**Figure S6.** Phylogeny construction using Parvovirus-GLUE

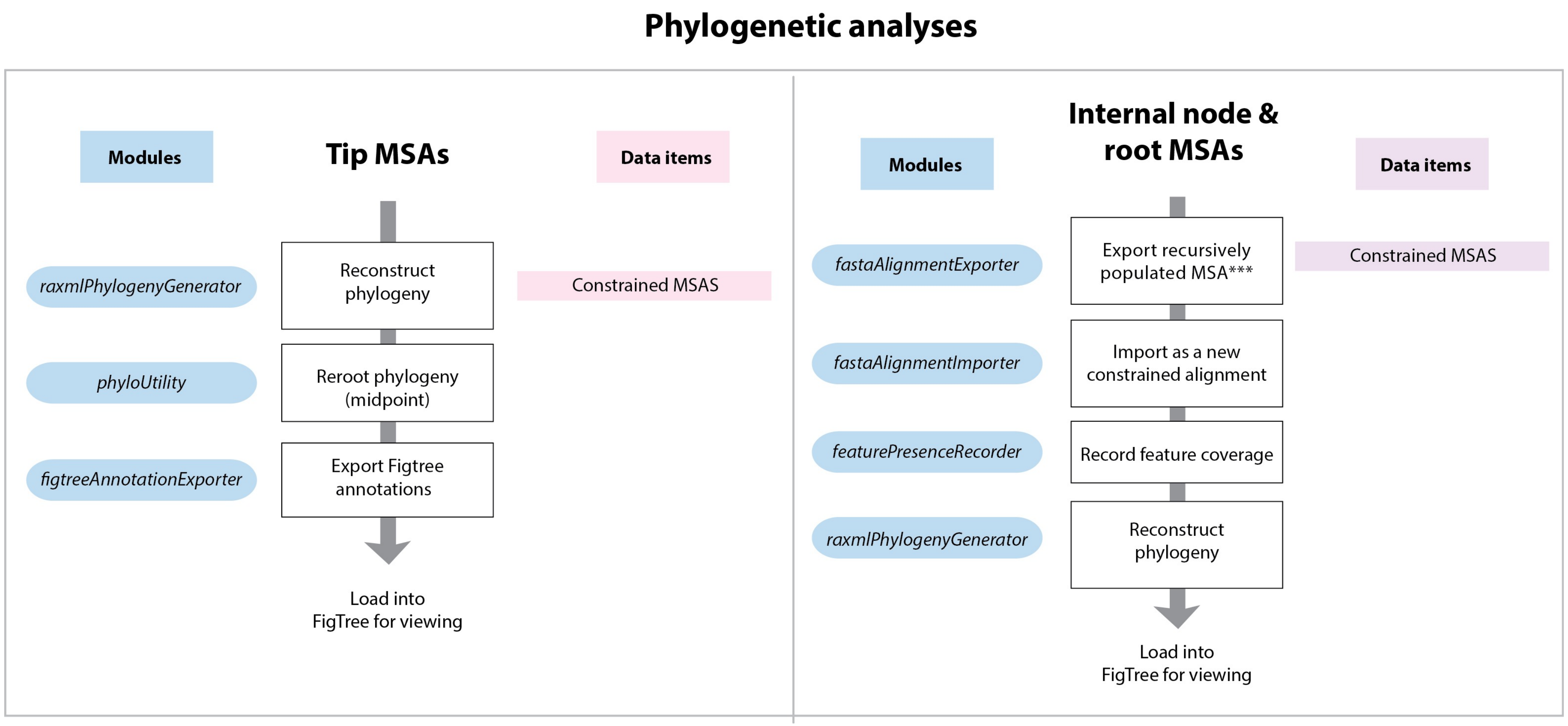

**Figure S7.** Comprehensive phylogenetic analysis of vertebrate parvoviruses (viruses only)

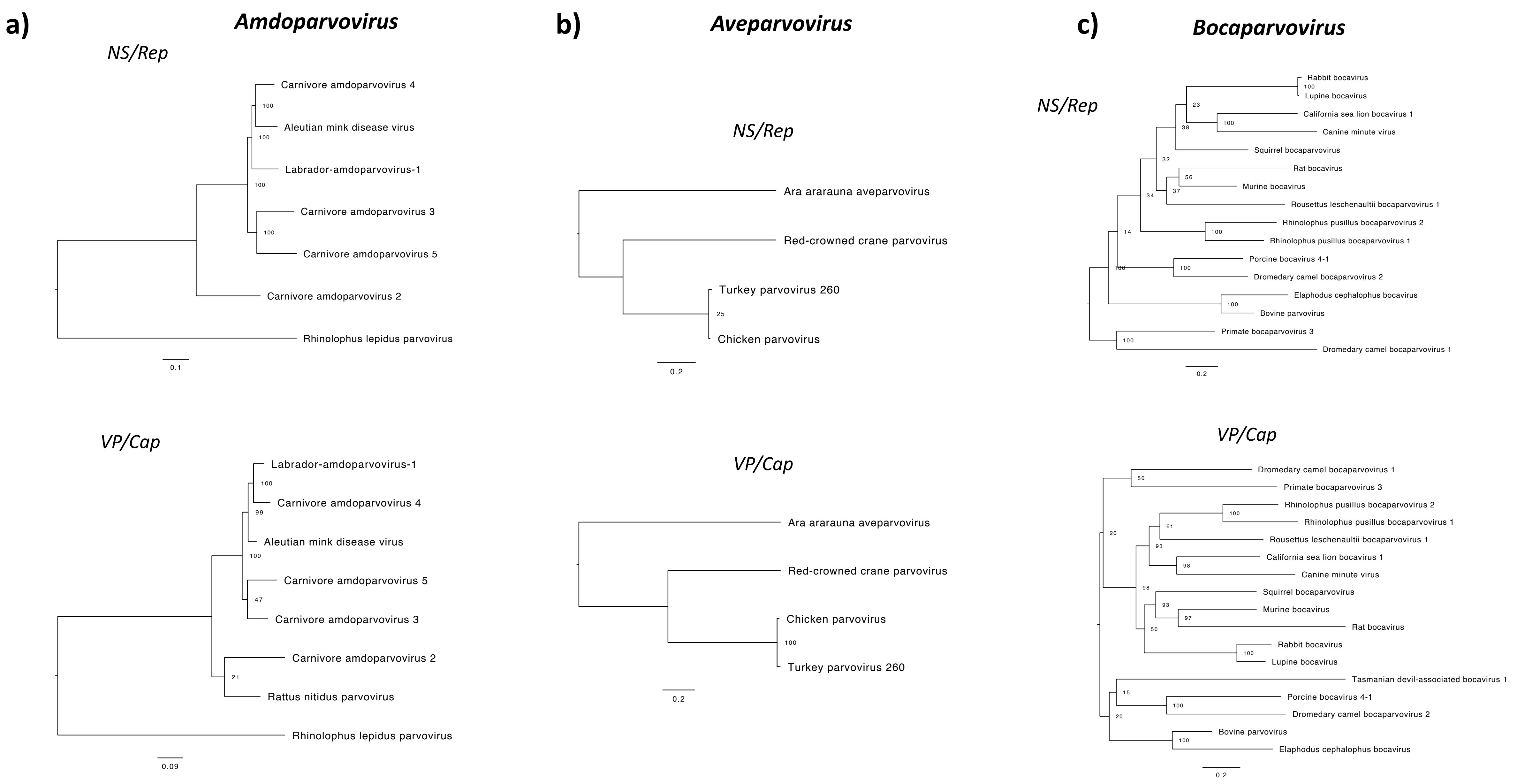

**Figure S7.** Comprehensive phylogenetic analysis of vertebrate parvoviruses (viruses only)

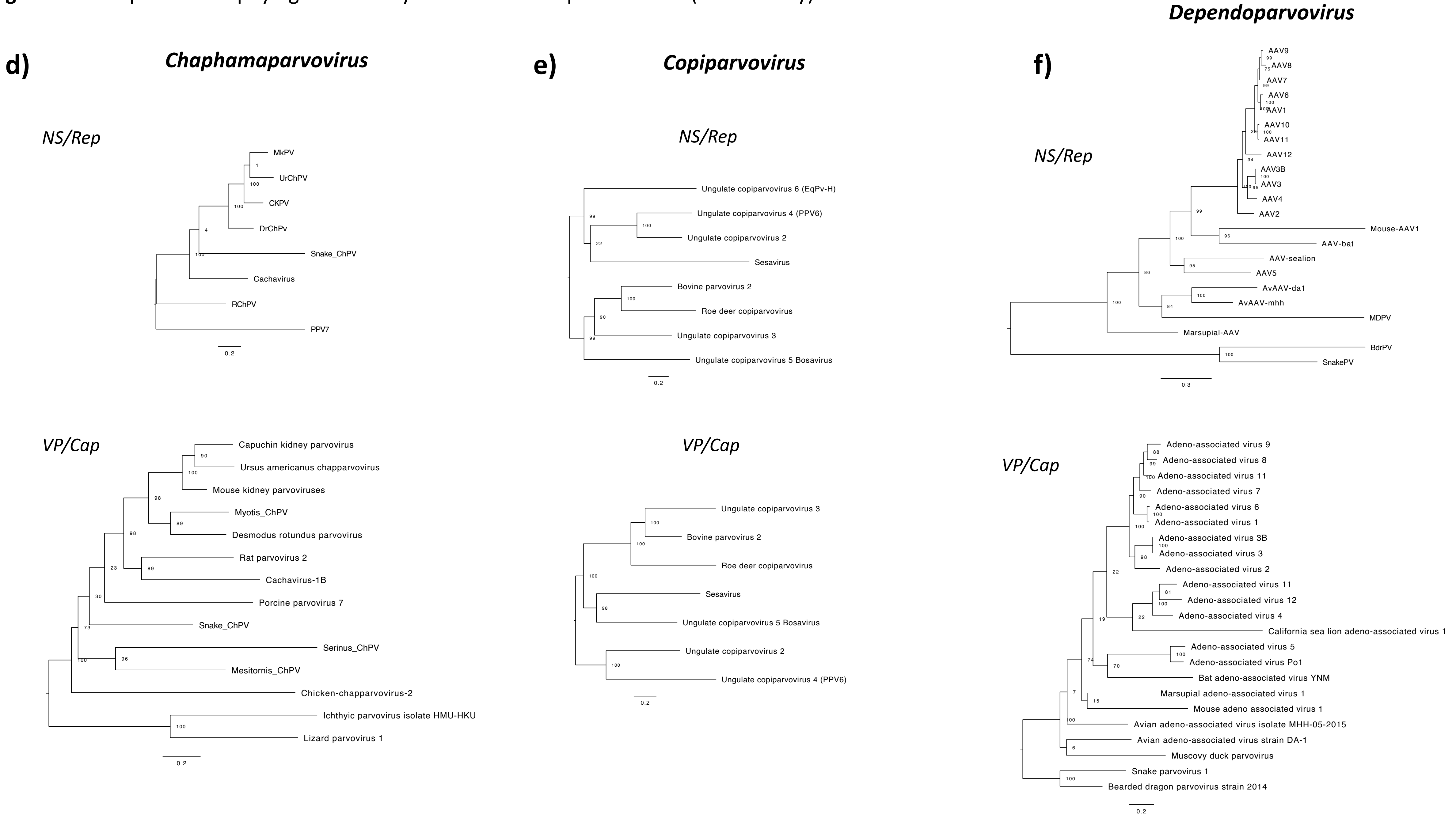

**Figure S7.** Comprehensive phylogenetic analysis of vertebrate parvoviruses (viruses only)

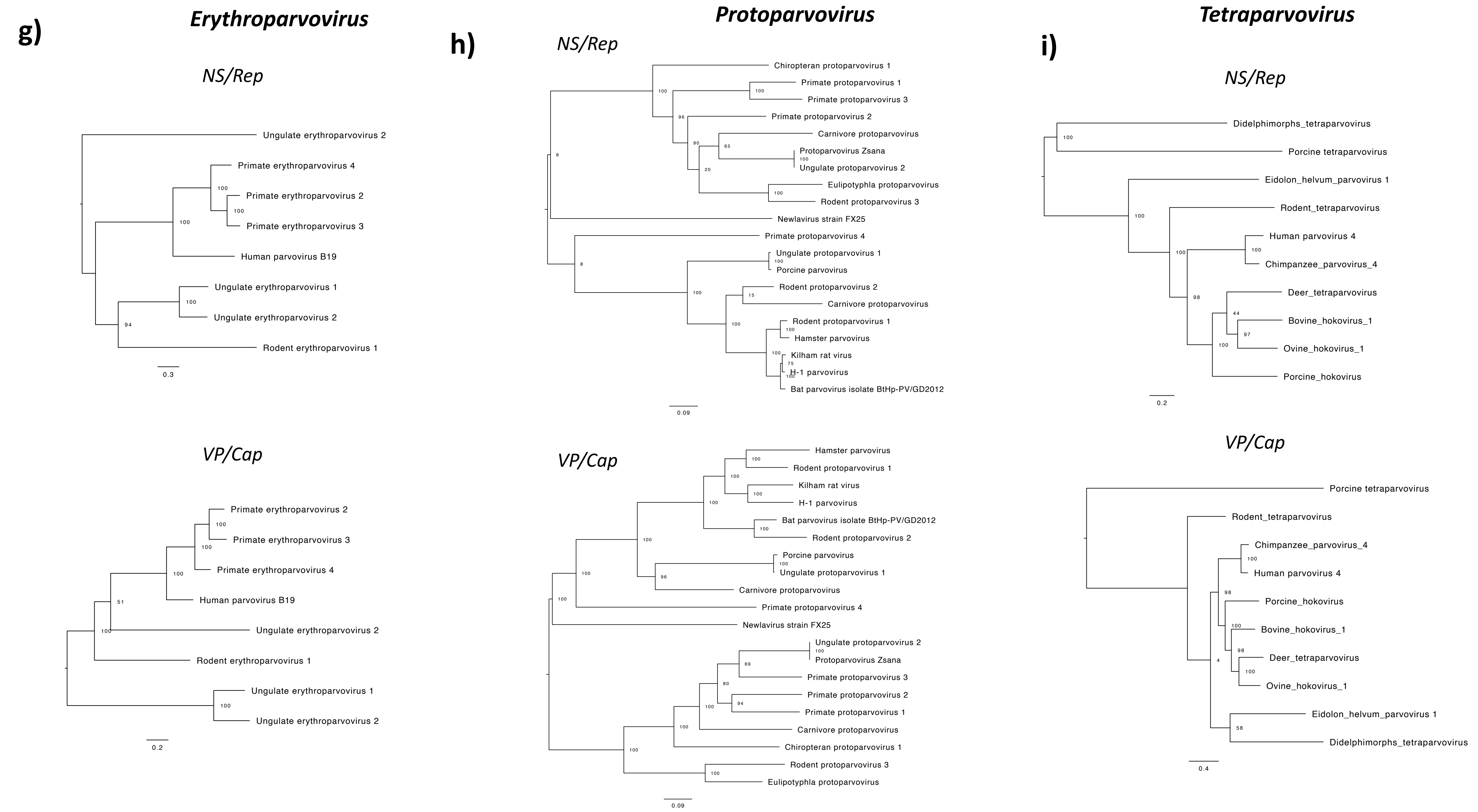

**Figure S8.** Comprehensive phylogenetic analysis of subfamily *Parvovirinae* including EPVs.

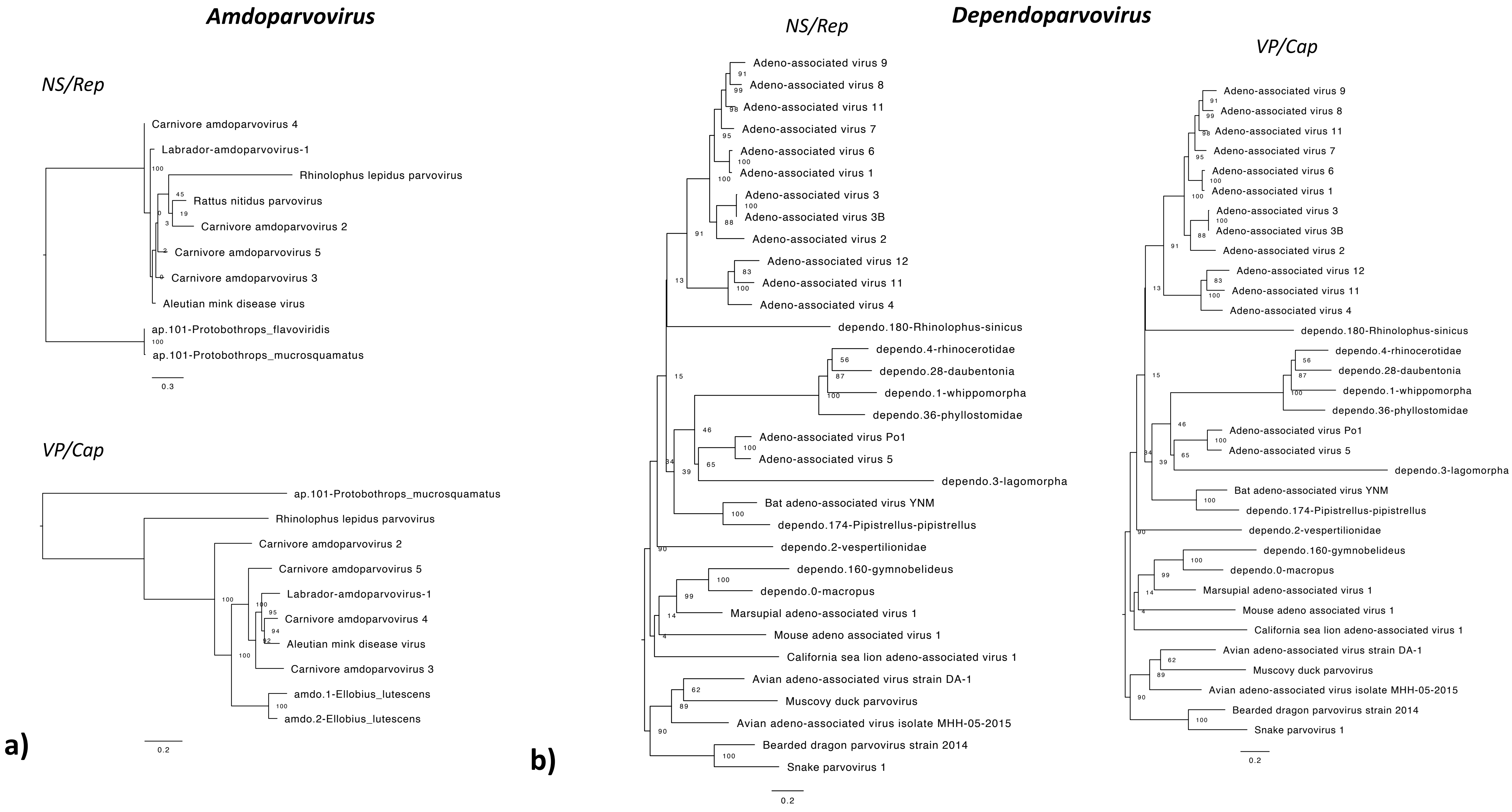

**Figure S8.** Comprehensive phylogenetic analysis of subfamily *Parvovirinae* including EPVs.

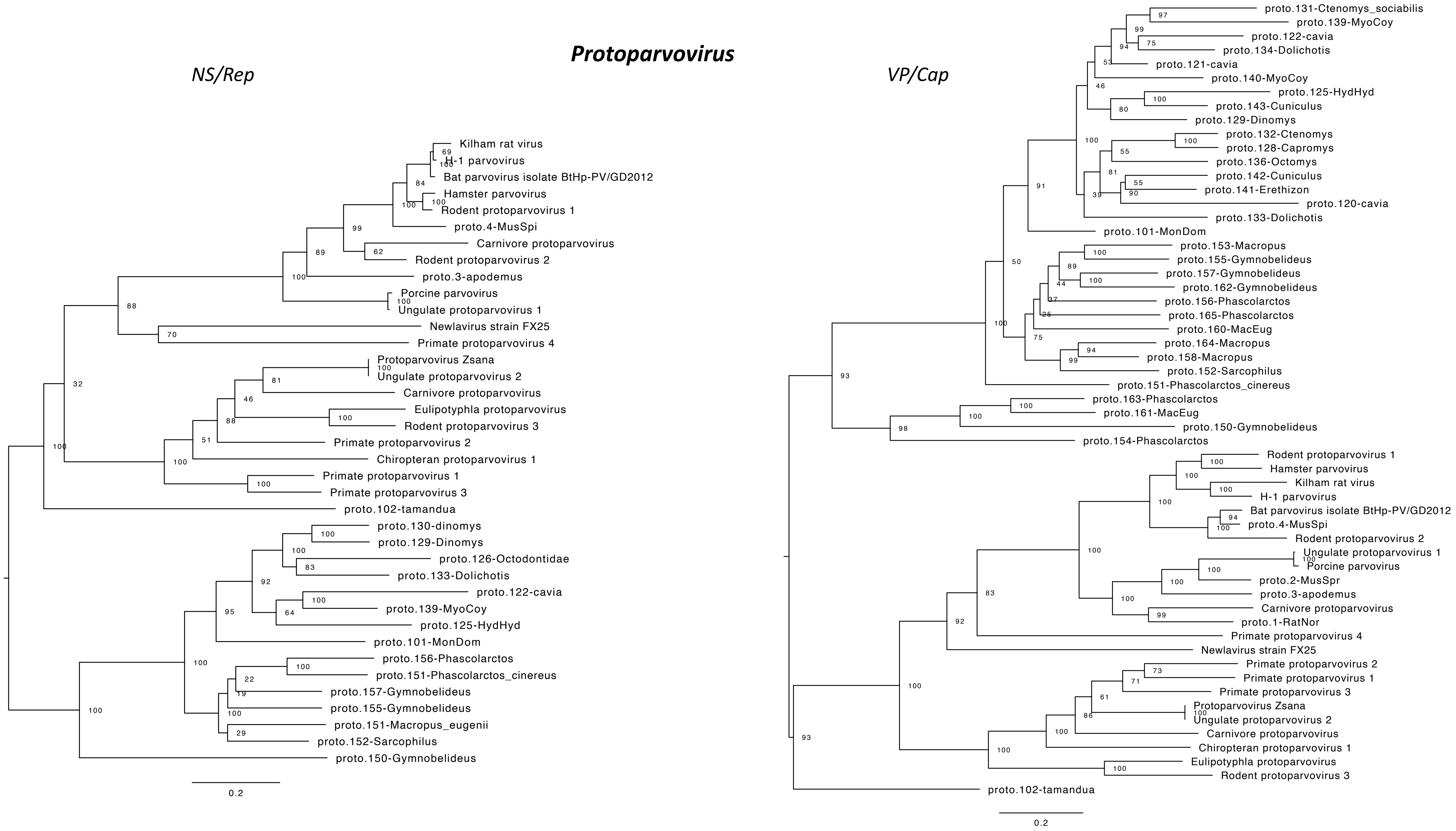


**c)**

**Figure S9.** Exogenous viruses identified in WGS and transcriptome databases.

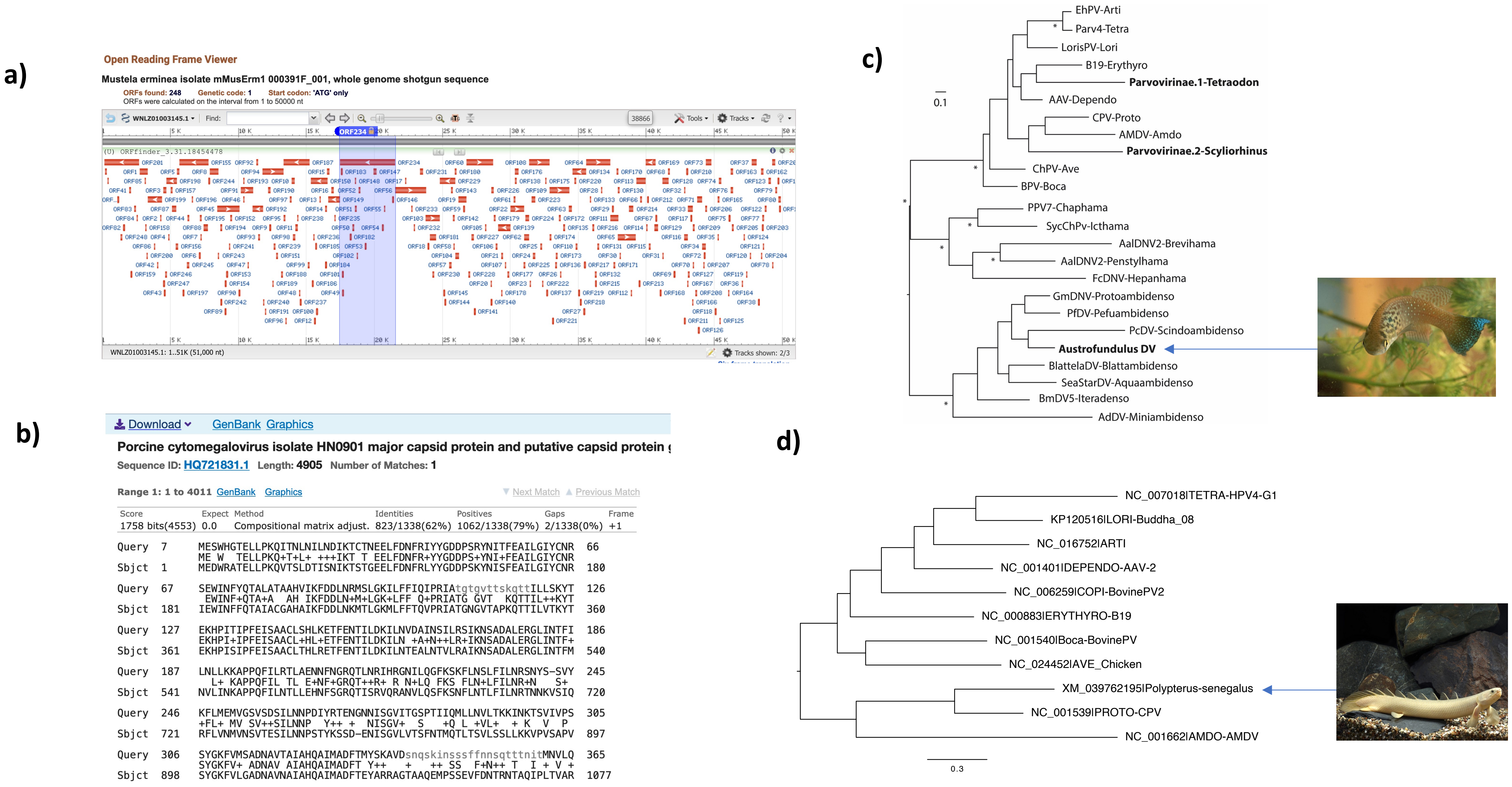

**Figure S10.** U94 alignment.

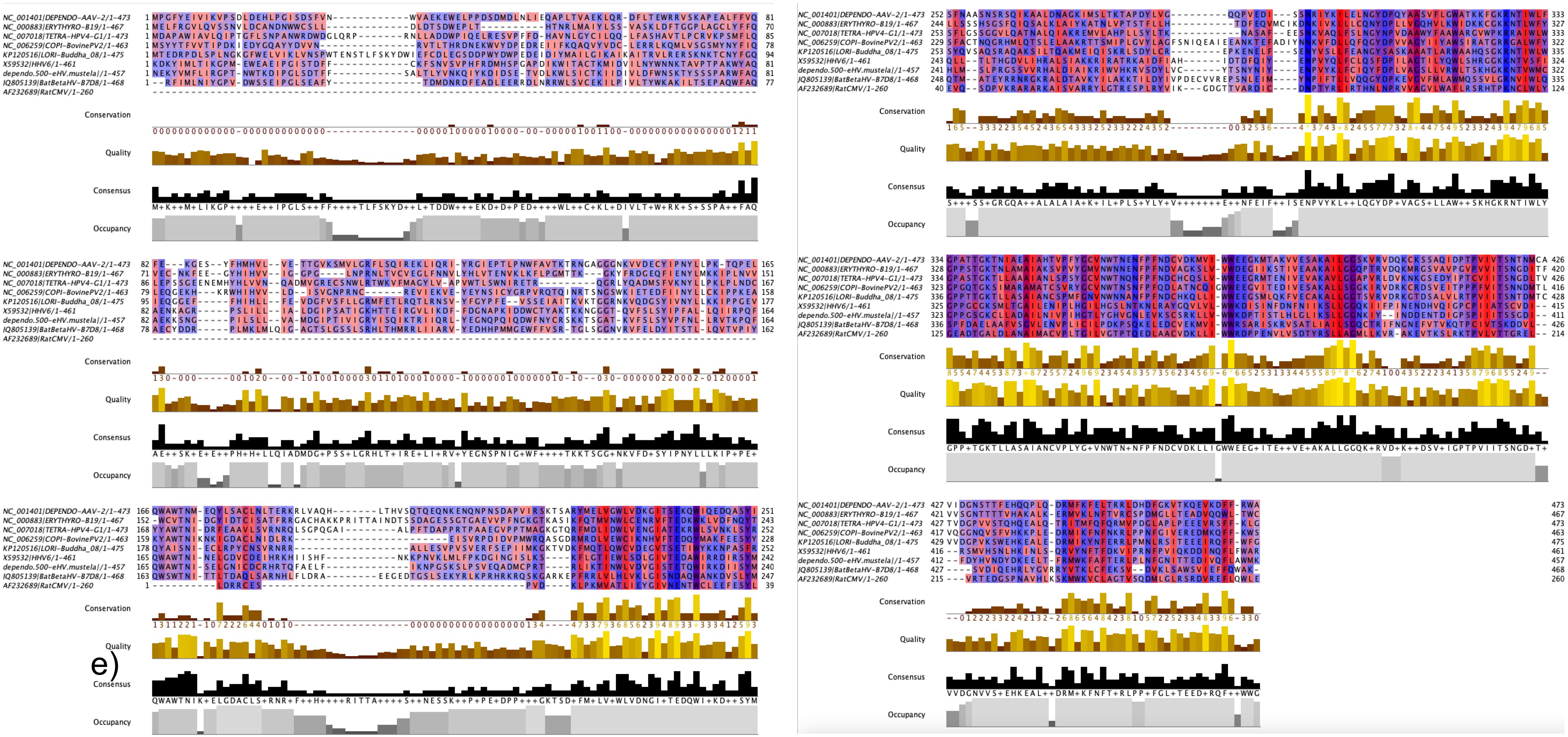

**Figure S11.** Evolution of protoparvoviruses.

a)

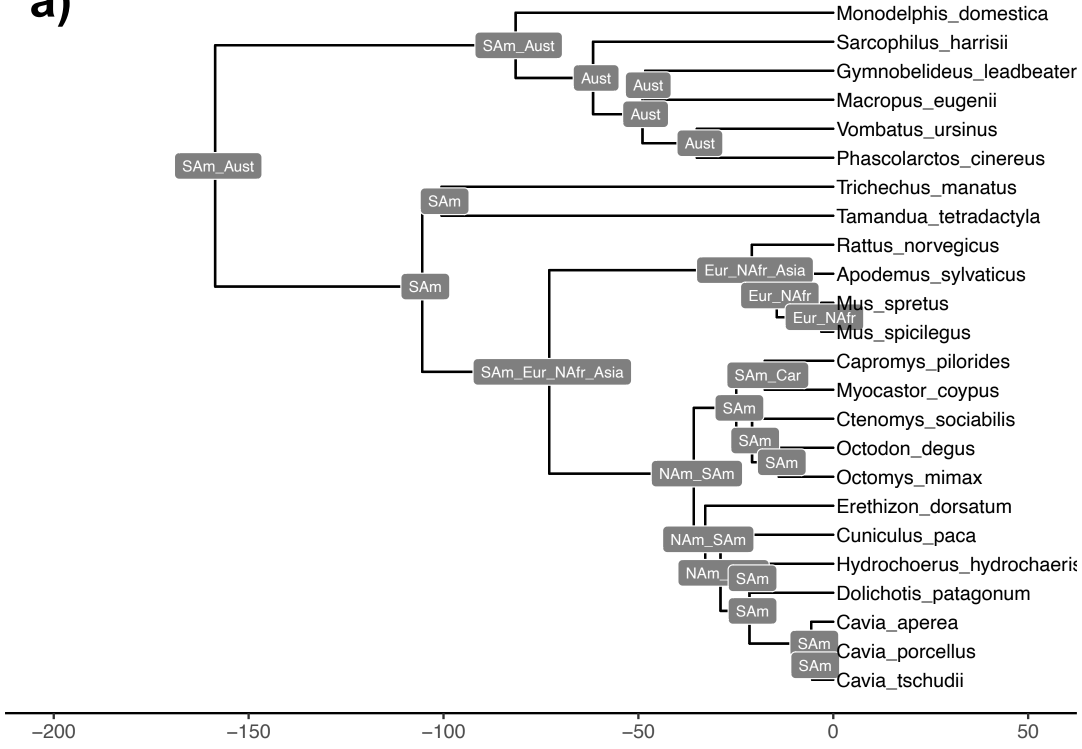


b)

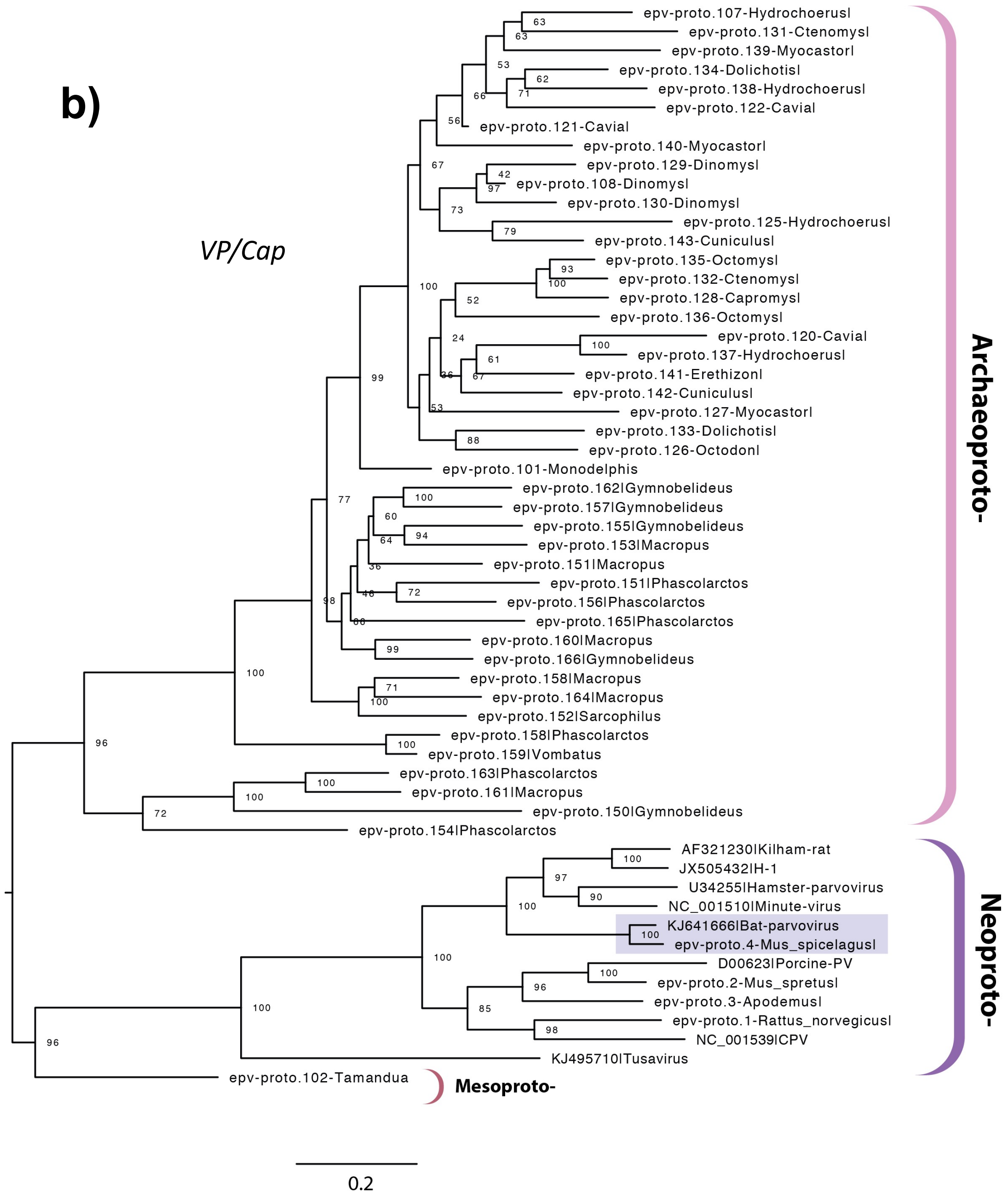

**Figure S12.** Dependoparvovirus VP/capsid phylogeny.

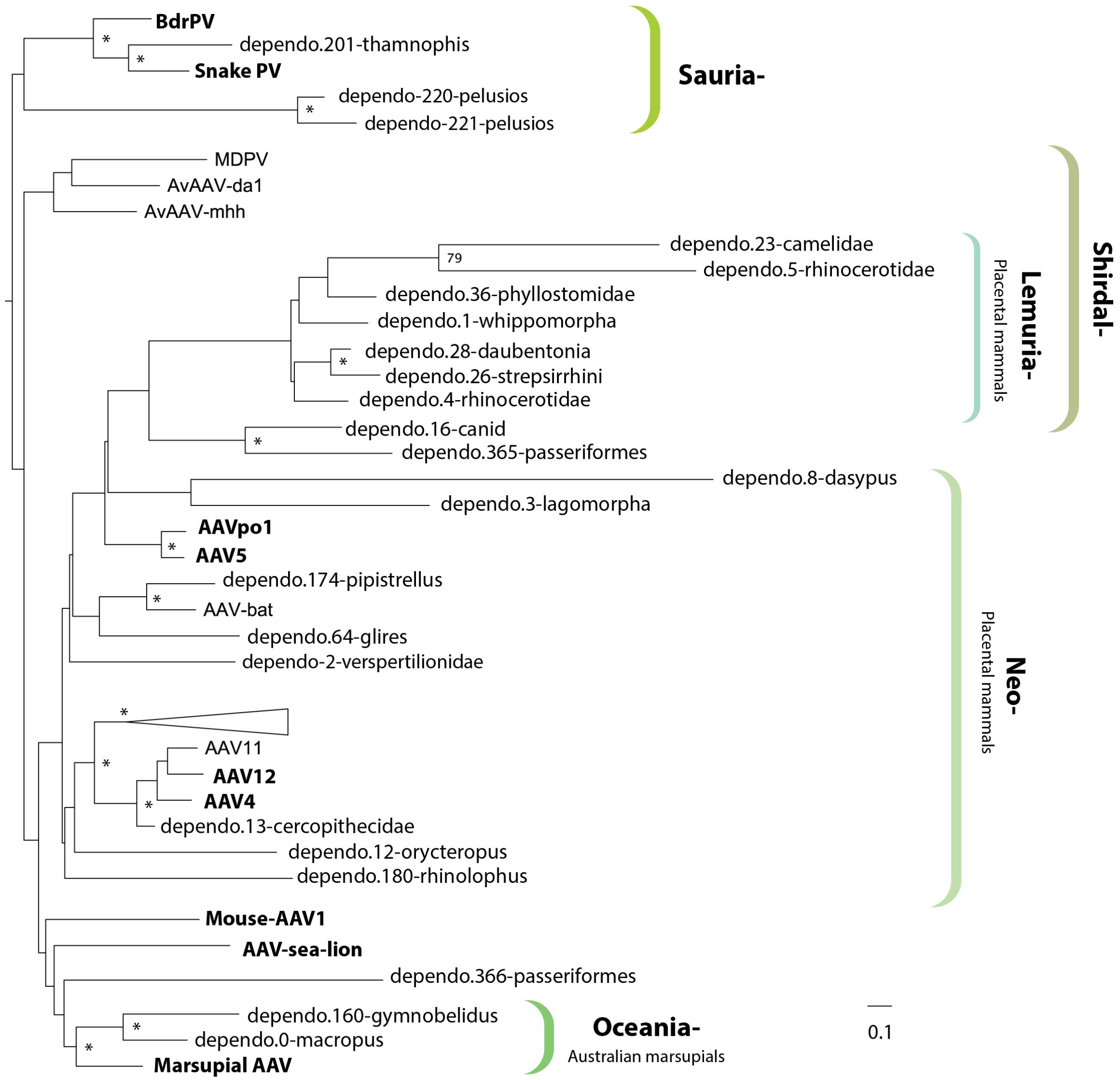

**Figure S13.** Multiple sequence alignment of dependoparvovirus-derived EPVs

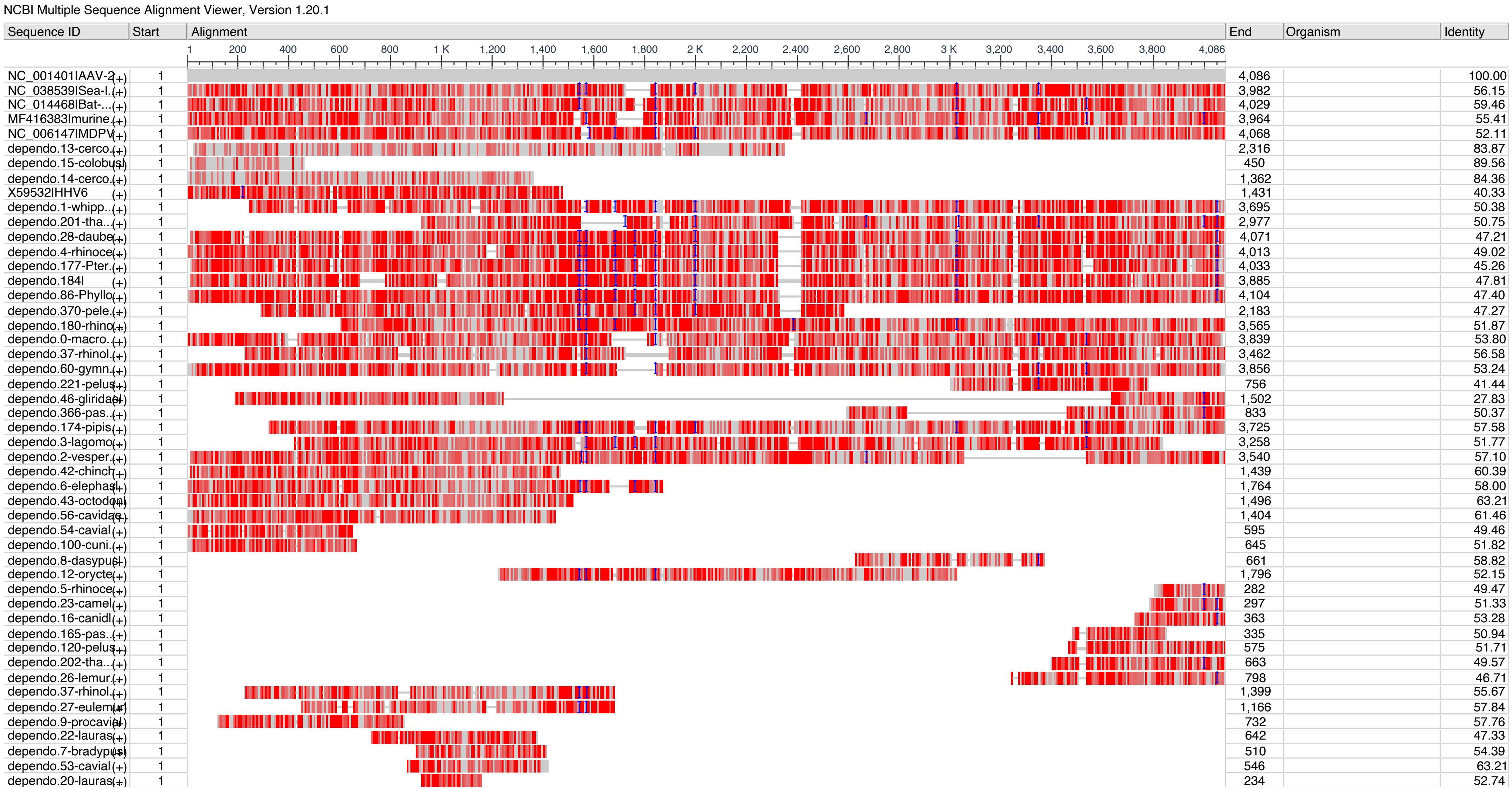

**Figure S14.** Evolution of the Erythroparvovirus genus.

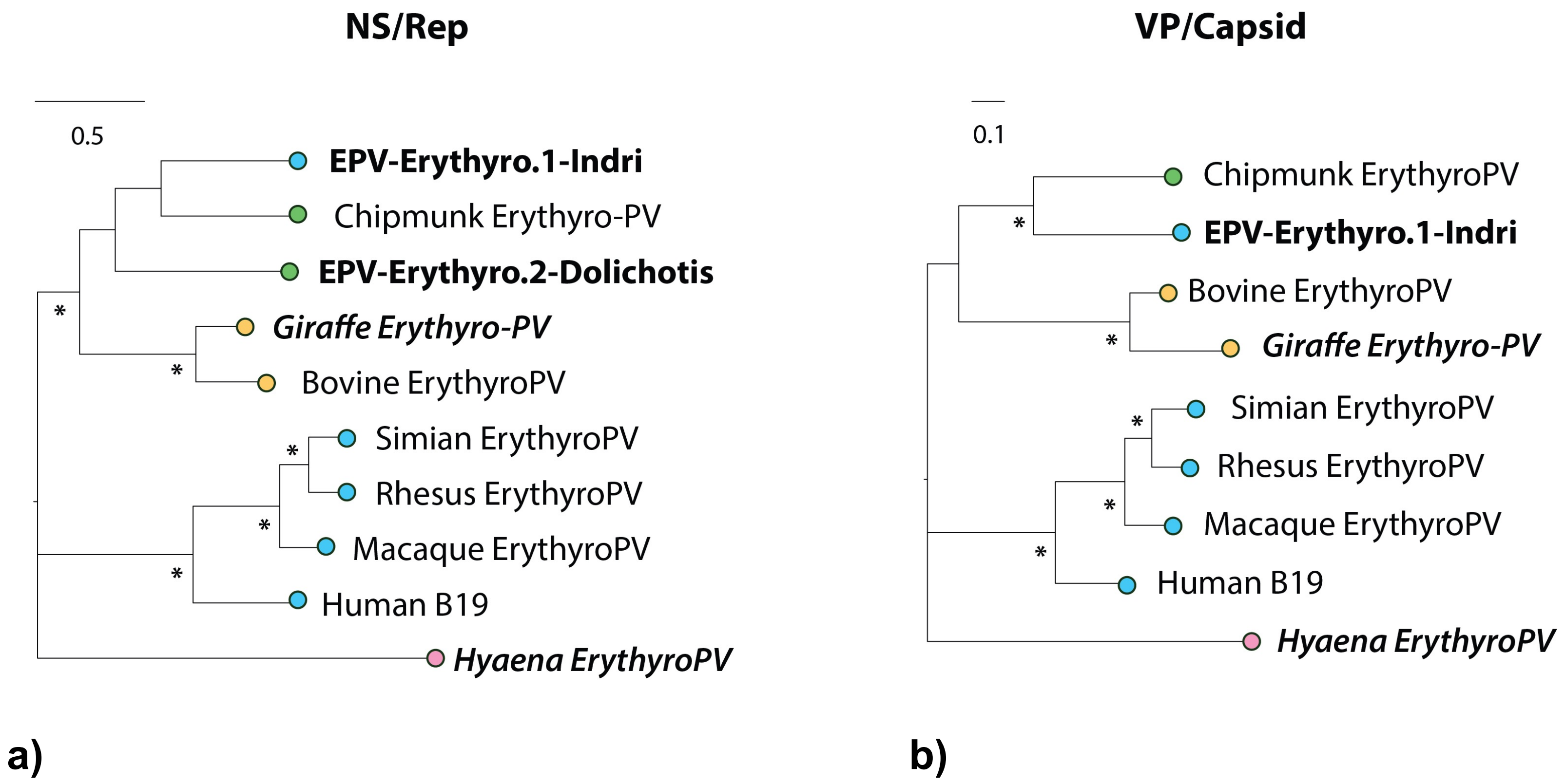

**Figure S15.** Evolutionary relationships of vertebrate hamaparvoviruses.

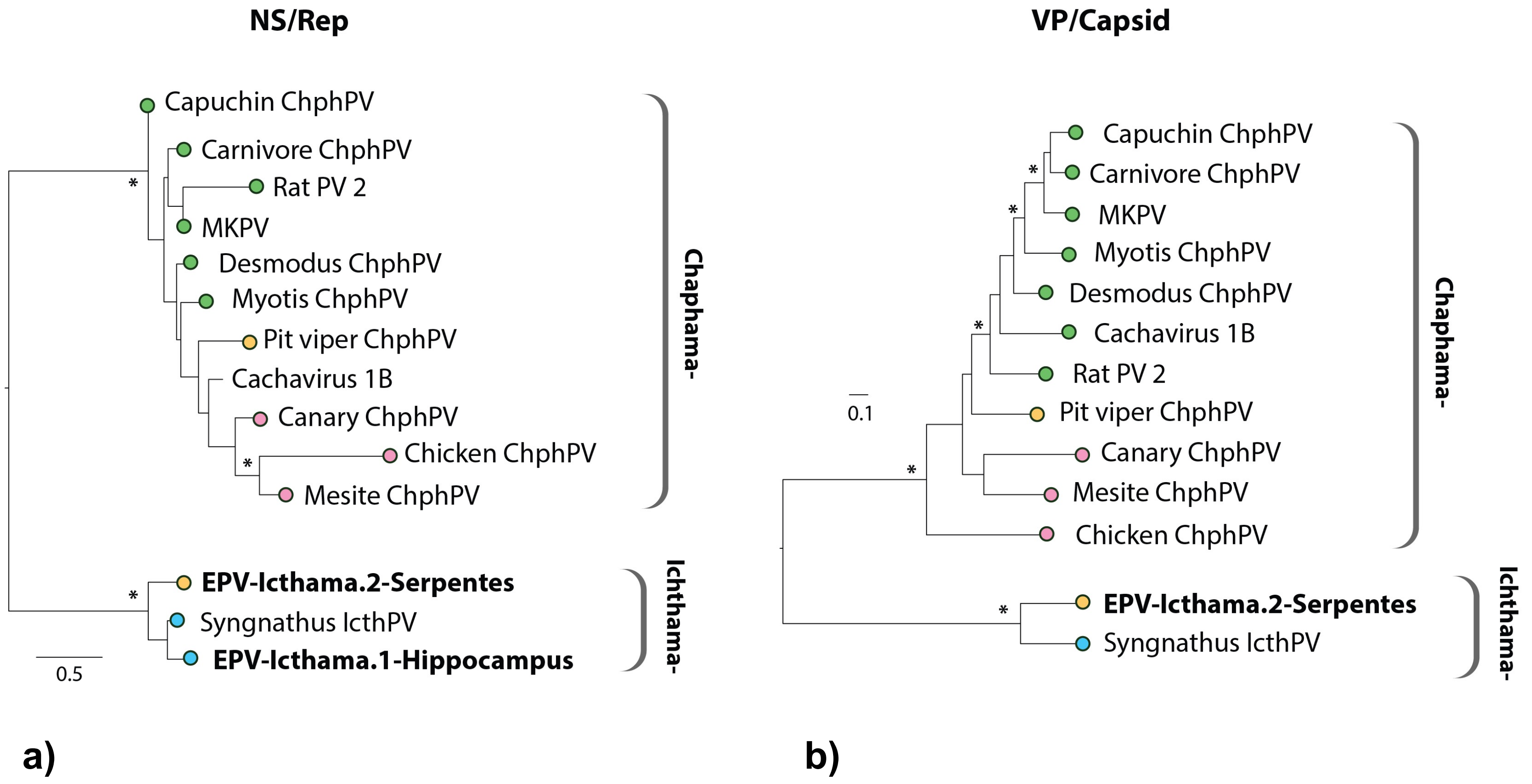
