## Supplemental figure legends for "Comparative analysis reveals the long-term co-evolutionary history of parvoviruses and vertebrates"

**Supplementary Figure Legends**

**Figure S1. Parvovirus-GLUE – an open resource for reproducible comparative analysis of parvovirus genome data.**

**(a)** The GLUE software framework allows users to develop flexible ‘projects’ oriented around the data items involved in comparative sequence analysis. Typically, these will include molecular sequence data, genome annotations, multiple sequence alignments (MSAs) and phylogenies, as well as diverse other forms of data. Loading projects into the GLUE ‘engine’ creates a relational database that captures the semantic relationships between data items. The database is constructed using GLUE’s native command layer, which can also be used to interact with the database and with commonly used bioinformatics programs (e.g., BLAST, RAxML, MAAFT). Working within this framework makes analyses reproducible and enables re-use and development of complex data items. **(b)** Loading projects into the GLUE ‘engine’ creates a relational database that captures the semantic relationships between data items. The database is constructed using GLUE’s native command layer, which can also be used to develop analysis protocols that utilise data items by interacting with the database and with commonly used bioinformatics programs (e.g., BLAST, RAxML, MAAFT) (see **Fig S2**). **(c)** Hosting of GLUE projects in an online version control system (e.g., GitHub) allows controlled collaborative development.

**Figure S2. The Parvovirus-GLUE resource build process.**

Flowchart showing the process through which (i) the core project database is constructed in the Parvovirus-GLUE resource (left); (ii) the Parvovirus-GLUE project database is extended via incorporation of genus-level project layers; (iii) the Parvovirus-GLUE project database is extended through the addition of EPV-specific project layers for individual parvovirus genera.

**Figure S3. A constrained alignment tree for the *Parvoviridae*.**

The schematic shows a represention of the constrained alignment tree data structure implemented in the *Parvoviridae*-GLUE resource. It comprises a set of multiple sequence alignments (MSAs) that are hierarchically linked to reflect taxonomic relationships and represents the entire *Parvoviridae* family (monotypic genera are not shown). Numbers on nodes correspond to the MSA numbers shown in **Table 1**. The dashed line illustrates how the project master reference sequence, carnivore protoparvovirus 1 (CPV-1), is present in the family, subfamily, and cross-genus MSAs, as well in the genus-level MSA for the protoparvoviruses, and thus links MSAs from root to tip. Moreover, since the relationships of all *Parvovirinae* reference sequences to CPV are captured - either directly or indirectly - by this MSA set, all of the MSAs are linked. This approach effectively allows a single underlying MSA to be used to perform phylogenetic reconstructions across a range of taxonomic levels within the family. Furthermore, because each MSA is constrained to a master reference, a standardised genomic coordinate space is imposed on all parvovirus species genomes.

**Figure S4. Compositional biases in parvovirus genomes.**

Heatmaps showing **(a)** nucleotide, **(b)** dinucleotide, and **(c)** amino acid composition of distinct genome features in reference parvoviruses. The columns (*x* – axis) are percent composition and the rows (*y* – axis) represent master references for *Parvovirinae* genera in *Parvoviridae*-GLUE. A colour key for infrequent (red) to more common (yellow) residues is shown with a histogram of the values shown in the heatmap.

**Figure S5.** **Genome screening *in silico*.**

The figure shows a schematic representation of the database-integrated genome-screening (DIGS) process used to identify EPV sequences.  **(1)** We collated a ‘reference sequence library’ representing all known parvovirus species and all known EPVs, and corresponding open reading frame (ORF) annotations. We used GLUE to derive a library of translated ORF sequences from these data and used these as ‘queries’ in tBLASTn-based searches of vertebrate WGS databanks. Hits were extracted to a relational database (DB). **(2)** Hits were initially classified by tBLASTx-based comparison to the translated ORF library, and these classifications were recorded, along with other information about the hit (species genome, and assembly version, hit sequence, coordinates and orientation) in a relational database. Hits that were within 1000bp of one another were concatenated and classified the via BLASTn-based comparison to a nucleotide-level reference library containing virus genomes and previously characterised EPVs. As we progressively identified novel EPV loci we incorporated them into this library. **(3)** Database-assisted analysis of similarity scores – combined with *ad hoc* phylogenetic anayses – were used to filter hits and identify sets of orthologous EPV insertions.

**Figure S6. Phylogeny construction using Parvovirus-GLUE.**

Flowcharts showing the process through which maximum likelihood phylogenies of endogenous parvoviral elements (EPVs) were constructed using Parvovirus-GLUE.

**Figure S7. Comprehensive phylogenetic analysis of vertebrate parvoviruses (viruses only).**

Panels (a-g) show maximum likelihood phylogenies showing evolutionary relationships within parvovirus genera infecting vertebrates. Phylogenies were generated from codon-level multiple sequence alignments of NS/Rep and VP/Cap gene sequences, as follows: (a) *Amdoparvovirus*; (b) *Aveparvovirus*; (c) *Bocaparvovirus*; (d) *Chaphamaparvovirus*; (e) *Copiparvovirus*; (f) *Dependoparvovirus*; (g) *Erythroparvovirus*; (g) *Protoparvovirus*; (h) *Tetraparvovirus*. Trees for individual EPV loci are available in the Parvovirus-GLUE online resource {Gifford, 2021 #237}.

**Figure S8. Comprehensive phylogenetic analysis of subfamily *Parvovirinae* including EPVs.**

Panels (a-g) display maximum likelihood phylogenies showing evolutionary relationships within parvovirus genera infecting vertebrates. Phylogenies were generated from codon- level multiple sequence alignments of NS/Rep and VP/Cap gene sequences, as follows: (a) *Amdoparvovirus*; (b) *Dependoparvovirus*; (c) *Protoparvovirus*.

**Figure S9. Exogenous viruses identified in WGS and transcriptome databases.**

**(a)** Open reading frame (ORF) prediction within the first 50 kilobases (kb) of a contig in WGS data for the stoat (*Mustela ermina*) that represents a putative betaherpesvirus, using NCBI’s ‘ORFfinder’ program. The polypeptide sequence encoded by the longest ORF (highlighted) was used as a BLAST ‘query’ in searches of NCBI nucleotide databases, and the most closely related sequence was found to be the polymerase gene porcine cytomegalovirus. Panel **(b)** shows the ‘alignment’ produced by this BLAST query. Panel **(c)** shows a phylogeny of the family *Parvoviridae* based on an alignment of the tripartite helicase domain and showing evolutionary relationships at the genus level*.* The position of a densovirus-like sequence recovered from WGS data of the annual killifish (*Austrofundulus* *truncatus*) is indicated. Panel **(d)** shows a phylogeny of the subfamily *Parvovirinae* based on an alignment of the NS protein. The position of a protoparvovirus-like sequence recovered from transriptome data obtained for the Senegalese bichir is indicated.

**Figure S10. U94 alignment.**

A polypeptide-level alignment of betaherpesvirus U94 genes. The alignment was constructed using MUSCLE and was imported into JalView for display purposes. The ‘hydrophobicity’ colour index was used to indicate conservation level.

**Figure S11. Evolution of protoparvoviruses.**

Panel **(a).** We examined the biogeography of *Protoparvovirus* through inferring the ancestral distributions of host species. A time-calibrated phylogeny of species harbouring EPV sequences was obtained from TimeTree. We combined the time-calibrated phylogeny and present continental distribution of host organisms to model ancestral distributions as shown in the tree. The most likely ancestral distribution at the root (158.60 million years ago) was found to be South America/Australia, implying an origin in Pangea. Panel **(b)** shows a maximum likelihood phylogenetic tree, based on an alignment of VP/capsid polypeptide sequences (544 amino acid residues) and showing the reconstructed evolutionary relationships between protoparvoviruses and protoparvoviruses-derived EPVs. The phylogeny was reconstructed using RAXML and the LG substitution model. Scale bar shows evolutionary distance in substitutions per site. Asterisks indicate nodes with >70% bootstrap support (1000 replicates). The close relationship between a neoprotoparvovirus-derived EPV identified in this study (EPV-Proto.4-MusSpre) and a recently identified, bat-associated protoparvovirus (“Pomona roundleaf bat protoparvovirus”), is highlighted**.**

**Figure S12. Dependoparvovirus VP/capsid phylogeny.**

A maximum likelihood phylogenetic tree, based on an alignment of VP/capsid polypeptide sequences (195 amino acid residues) and showing the reconstructed evolutionary relationships between dependoparvoviruses and dependoparvoviruses-derived EPVs. The phylogeny was reconstructed using RaXML and the LG model of amino acid substitution.

**Figure S13. Multiple sequence alignments of EPV and virus sequences.**

Figure summarising dependoparvovirus EPV coverage relative to a master reference sequence (adeno-associated virus 2).

**Figure S14. Evolution of the *Erythroparvovirus* genus**

Maximum likelihood phylogenetic trees showing the reconstructed evolutionary relationships between erythroparvoviruses and erythroparvovirus-derived EPVs. The Rep/NS phylogeny (left) was constructed using a polypeptide-level multiple sequence alignment (MSA) spanning 500 amino acid residues (substitution model= LG likelihood). The VP/Capsid phylogeny (right) was constructed using a multiple sequence alignment (MSA) spanning 780 amino acid residues (substitution model= LG likelihood). Scale bars show evolutionary distance in substitutions per site. Asterisks indicate nodes with >70% bootstrap support (1000 replicates). Bold taxa labels indicate EPV taxa, newly characterised viral taxa are shown in bold italic text while previously characterised viral taxa are shown in regular text.

**Figure S15. Evolutionary relationships of vertebrate hamaparvoviruses.**

Maximum likelihood phylogenetic trees showing the reconstructed evolutionary relationships between contemporary vertebrate hamaparvoviruses and hamaparvovirus-derived EPVs. The Rep/NS phylogeny (left) was constructed using a multiple sequence alignment (MSA) spanning 580 amino acid residues (substitution model= LG likelihood). The VP/Capsid phylogeny (right) was constructed using a multiple sequence alignment (MSA) spanning 450 amino acid residues (substitution model= LG likelihood). Scale bars show evolutionary distance in substitutions per site. Asterisks indicate nodes with >70% bootstrap support (1000 replicates). Coloured circles indicate host associations as follows: blue=fish; pink=birds; yellow=reptiles; green=mammals. Bold taxa labels indicate EPV taxa, while viral taxa are shown in regular text.
